# Patch-Clamp Single-Cell Proteomics in Acute Brain Slices: A Framework for Recording, Retrieval, and Interpretation

**DOI:** 10.1101/2025.09.15.675920

**Authors:** Larry Rodriguez, Jolene Diedrich, Le Sun, Blake Tsu, Stefanie Kairs, Roman Vlkolinsky, Christopher A. Barnes, Aline M. A. Martins, Marisa Roberto, John R. Yates

## Abstract

Single-cell proteomics (SCP) is a powerful method for interrogating the molecular composition of neurons, yet its application to acute brain slices has remained limited. Patch-clamp electrophysiology provides direct information on neuronal excitability, synaptic inputs, and ion channel function, making it a natural partner for SCP. However, combining these techniques introduces unique challenges. For instance, after patch-clamping a neuron, its soma must be physically retrieved, and variability during extraction from the brain slice may influence how well proteomic measurements reflect *in situ* physiology. Here, we introduce a framework for contextualizing patch-SCP outcomes, with an emphasis on retrieval quality (material yield and soma-enriched synaptic content). We used an indiscriminate shotgun strategy in which all patched neurons were collected regardless of electrophysiological outcome to assess soma retrieval in an exploratory dataset of rat medial prefrontal cortex pyramidal neurons. Capacitance during gigaseal-preserved retrieval correlated with protein identifications, suggesting that proteome yield could be linked to soma size. Preservation of neuronal spiking during relocation tended to be associated with broader synaptic enrichment and recovery of transmembrane proteins. By comparison, torn or aspirated neurons produced small proteomes with poor synaptic representation and neurons with little to no characterization displayed more variable outcomes. These results demonstrate that patch-SCP can be used to assess soma retrieval and they provide a framework for interpreting how electrophysiological context and soma retrieval quality shape single-neuron proteomic measurements in semi-intact circuits.

## Introduction

Patch-clamp electrophysiology performed in acute brain slices is the gold standard for assessing the functional properties of individual neurons. In the whole-cell configuration, a glass micropipette is used to establish electrical continuity with the intracellular space of a single neuron, which enables precise measurement of voltage- and ligand-gated ion channel activity, synaptic inputs, and intrinsic excitability. Advances in single-cell “omics” have extended the use of patch-clamp electrodes beyond the measurement of physiological responses to include the physical collection of neuronal material for molecular profiling. Early efforts combined patch-clamp electrophysiology with targeted transcript detection using single-cell RT-PCR, and were later extended to whole-transcriptome RNA sequencing in what became known as Patch-seq[1–5]. More recently, progress in mass spectrometry-based single-cell proteomics (SCP) has made it possible to identify proteins from a neuron that was patch-clamped (i.e. patch-SCP). Initially, patch-SCP in intact brain slices relied on aspirating cytoplasmic contents through the recording electrode [6–8]. Subsequent work extended the approach to human iPSC-derived neurons, where whole somas could be collected and analyzed [9]. While these advances illustrate the versatility of patch-SCP, the process for coordinating a neuron’s electrophysiology with its protein composition remains challenging.

Electrophysiologists typically study ion channels, G protein-coupled receptors (GPCRs), and synapse-associated proteins, which are often low-abundance and membrane-embedded, making them difficult to detect by mass spectrometry. Thus, to expand the detailed neurophysiological insight gained from acute brain slice recordings, SCP workflows must be able to detect proteins that contribute to the electrical properties of the neuron (e.g. excitability), which may be localized at distal compartments rather than in the soma. This means that even when patch-clamp reveals distinct neurophysiological properties, the mass spectra may not capture the molecules that affect those signals. For example, patch-SCP in the locus coeruleus of mice revealed sex-specific differences in both the proteomes and intrinsic excitability of noradrenergic neurons, although collection and analysis were limited to the cytoplasm, leaving the molecular drivers of these differences difficult to identify[8]. Similarly, a patch-SCP platform applied to Alzheimer’s disease hiPSC-derived neurons found an association between protein expression and a hyperexcitable phenotype, yet a lack of compartment-specific synaptic and membrane proteins raised the question of whether local synaptic activity could be adequately captured by the platform[9].

These studies demonstrate that to uncover the molecules regulating neuronal physiology, patch-SCP must overcome several technical bottlenecks, such as sampling depth, compartmental bias, and the mechanical challenges of retrieving neurons from brain slices. Mechanical retrieval itself represents a fundamental constraint; proteomic recovery can vary substantially if the soma tears during withdrawal or if significant material is lost, which can occur during sample collection and/or preparation. To mitigate these challenges, many patch-SCP studies have adopted an “all-or-nothing” strategy, in which only neurons with stable recordings are targeted for collection and only samples exceeding predefined protein-identification thresholds are analyzed[8, 9]. While this approach can be used to enrich for high-quality datasets, it also assumes that electrophysiological recordings reliably predict proteomic recovery, an assumption that has not been systematically tested. Because manual patch-clamp recordings are labor-intensive, patch-SCP workflows benefit from analytical frameworks that maximize interpretability across all retrieval outcomes. Such approaches enable systematic evaluation of retrieval quality, sample handling, and proteomic yield, rather than discarding partial or unsuccessful collections *a priori*.

Here, we introduce a framework for aligning patch-clamp electrophysiology with MS-based single-cell proteomics by explicitly accounting for variability in soma retrieval. To implement this framework, we developed a shotgun patch-SCP strategy. This approach systematically captures electrophysiological and retrieval outcomes to examine somatic protein recovery from single neurons in acute rat brain slices. By extending core electrophysiological principles to proteomic sampling and maintaining the gigaseal during soma retrieval, we explored the relationship between neuronal size and proteome yield. Importantly, neurons were collected for proteomic analysis regardless of whether the high-resistance seal was preserved, lost, or never established. This indiscriminate collection strategy enabled us to evaluate how retrieval integrity influences proteomic depth and composition across the dataset. We applied this framework to Layer 2/3 pyramidal neurons in the rat medial prefrontal cortex (mPFC), a population chosen for its relatively large soma, its rich synaptic proteome, and relevance to neuropsychiatric disorders[10, 11]. Using patch-SCP, we detected thousands of proteins from single neuronal soma, including ion channels, GPCRs, and synaptic proteins that govern neuronal properties. Moreover, we demonstrated how partially compromised retrievals can be leveraged to define contextual limitations and interpret variability in SCP outcomes. Collectively, this work illustrates how patch-SCP can be used to analyze semi-intact neural circuits.

## Results and Discussion

### A Framework for Interpreting Patch-Clamp SCP Recordings

Whole-cell patch-clamp in acute brain slices is a powerful approach for investigating neuronal physiology because it provides direct access to the currents and potentials that govern excitability. After forming a high resistance “gigaseal” between the micropipette and soma, a patch of the neuron’s membrane is ruptured, which makes the electrode electrically continuous with the intracellular space (Figure S1). Current-clamp configuration allows measurement of intrinsic membrane properties such as capacitance (C, which is proportional to surface area or size of the neuronal soma) and resistance (R_M_, which reflects how effectively ionic current flow is restricted across the membrane and can be used as a proxy for neuronal integrity). The active membrane properties of the neuron (mediated by voltage-gated sodium, potassium, and calcium channels) can also be assessed, including the generation and pattern of action potentials. Voltage-clamp configuration can measure excitatory and inhibitory postsynaptic currents which reflect connectivity and ongoing network activity in the slice. Neuronal physiology *in situ* can be deconstructed into ion channel and synaptic activity using this “patch-clamp circuit”. When the gigaseal is intact, the circuit and its features can be examined in a stable and meaningful manner; without it, recordings are not possible due to electrical discontinuity with the soma.

With these principles in mind, the rationale for combining SCP with patch-clamp is to provide molecular context for the electrophysiological properties of individual neurons. However, because the patch-clamp technique is performed manually, exploratory studies must be carried out on a small scale. With limited resources, a critical consideration for performing patch-SCP in acute brain slices should be the process of neuron retrieval or extraction. Patch-SCP isolates the patched soma and a stochastic assortment of its immediately adherent structures (rather than the full neuronal arbor). Retrievals that are incomplete or inconsistent may cause protein loss that impedes the mass spectrometry detection of proteins involved in the electrophysiological responses observed *in situ*. For example, a neuron may display sodium channel-dependent spiking, but if the axonal initiation segment (AIS) of the soma is only partially collected, then the Na_V_ subunits that mediated the response may be absent or below the limit of detection of the mass spectrometer. Soma-distal localization and/or inefficient proteomic workflows (i.e. under-recovery of hydrophobic proteins [12]) can cause “false negatives” which complicate patch-SCP interpretation, particularly for low-abundance transmembrane proteins.

Here, our goal was to assess the capabilities of our patch-SCP process. To accomplish this, we developed a framework for interpreting patch-SCP outcomes (Figure 1) by extending the core principles of patch-clamp electrophysiology to soma retrieval and incorporating methodological guidelines for SCP[13]. We reasoned that if we could preserve electrical continuity with the soma, then we might be able to reduce the uncertainty associated with losing synaptic structures during the retrieval process. This means maintaining the initial *in situ* whole-cell configuration via the gigaseal throughout the retrieval process, which provides insight into how the passive and active properties of the neuronal soma change in response to retrieval from the slice. However, the delicate nature of patch-clamping means that the most likely outcome when retrieving the soma is loss of the gigaseal. If the integrity of the soma at the time of recovery is uncertain, then the ability to associate an electrophysiological property (*in situ*) with the proteome retrieved will depend on the robustness of the analytical workflow (i.e. alignment of recovering proteins responsible for *in situ* responses). Alternatively, one may fail to form a gigaseal with the membrane, which provides no electrophysiological data and is often excluded from SCP analysis[8, 9]. Figure 1 summarizes what properties can be assessed under each patch-SCP configuration in our framework and the context each outcome provides.

**Figure 1.**
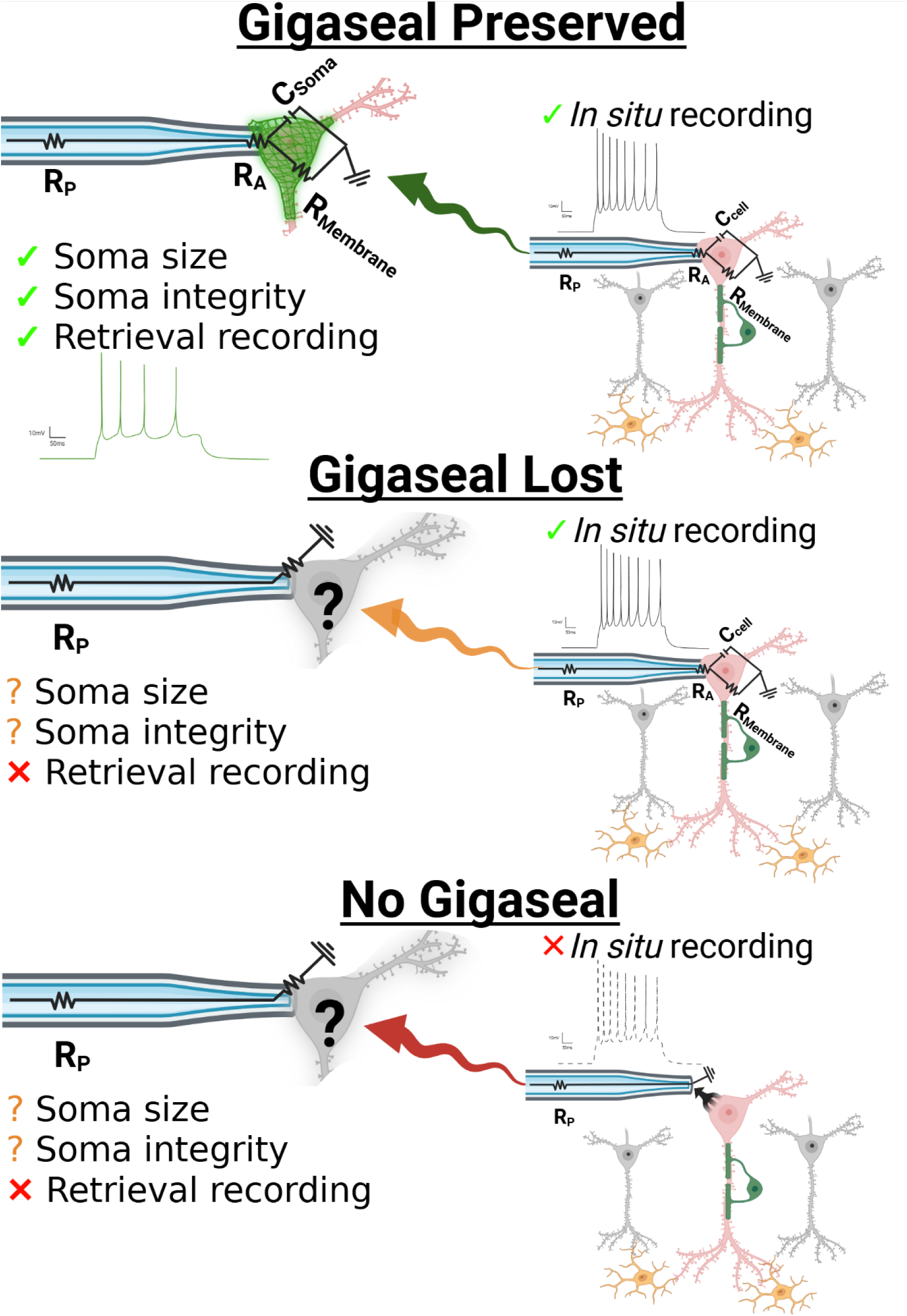
Conceptual framework for interpreting recordings from patch-SCP outcomes. Whole-cell patch-clamp provides access to information about a neuron’s intrinsic and synaptic properties, but retrieval introduces variability in how much of the soma (and its compartments) are recovered for proteomic analysis. Collection outcomes fall into three categories: (i) gigaseal preserved during retrieval, enabling continued electrophysiological monitoring of soma properties, (ii) gigaseal lost during retrieval, leaving pre-retrieval recordings valid but recovery uncertain, or (iii) no gigaseal, in which somas can still be collected but without functional characterization. Abbreviations: R_p_, pipette resistance; R_A_, access resistance; R_M_, membrane resistance; C_cell_, cell capacitance; C_Soma_, soma capacitance. Created with BioRender.

While recordings performed during retrieval might improve the interpretive confidence of patch-SCP (e.g. providing surrogate measures for how much neuronal material is collected or resolving the condition of the soma during the retrieval process), it does not guarantee preservation of all neuronal compartments. As shown in Figure 2, the more distal dendritic or axonal domains are likely to be retained in the slice because their physical connection is mediated by adhesion molecules[14]. This also means that if proximal synaptic connections can be collected with the soma, the postsynaptic terminals belonging to the soma will likely be isolated with some of its presynaptic partners. Given the uncertain nature of sample loss, we reasoned that soma collections lacking electrophysiological recordings could still be useful for assessing 1) the consistency of how we retrieve soma from the tissue, 2) proteome recovery, 3) MS performance, and 4) variability in our workflows. Considering that our goal was to explore our patch-SCP capabilities efficiently, we did not impose selection or exclusion criteria for either soma retrieval or MS analysis.

**Figure 2.**
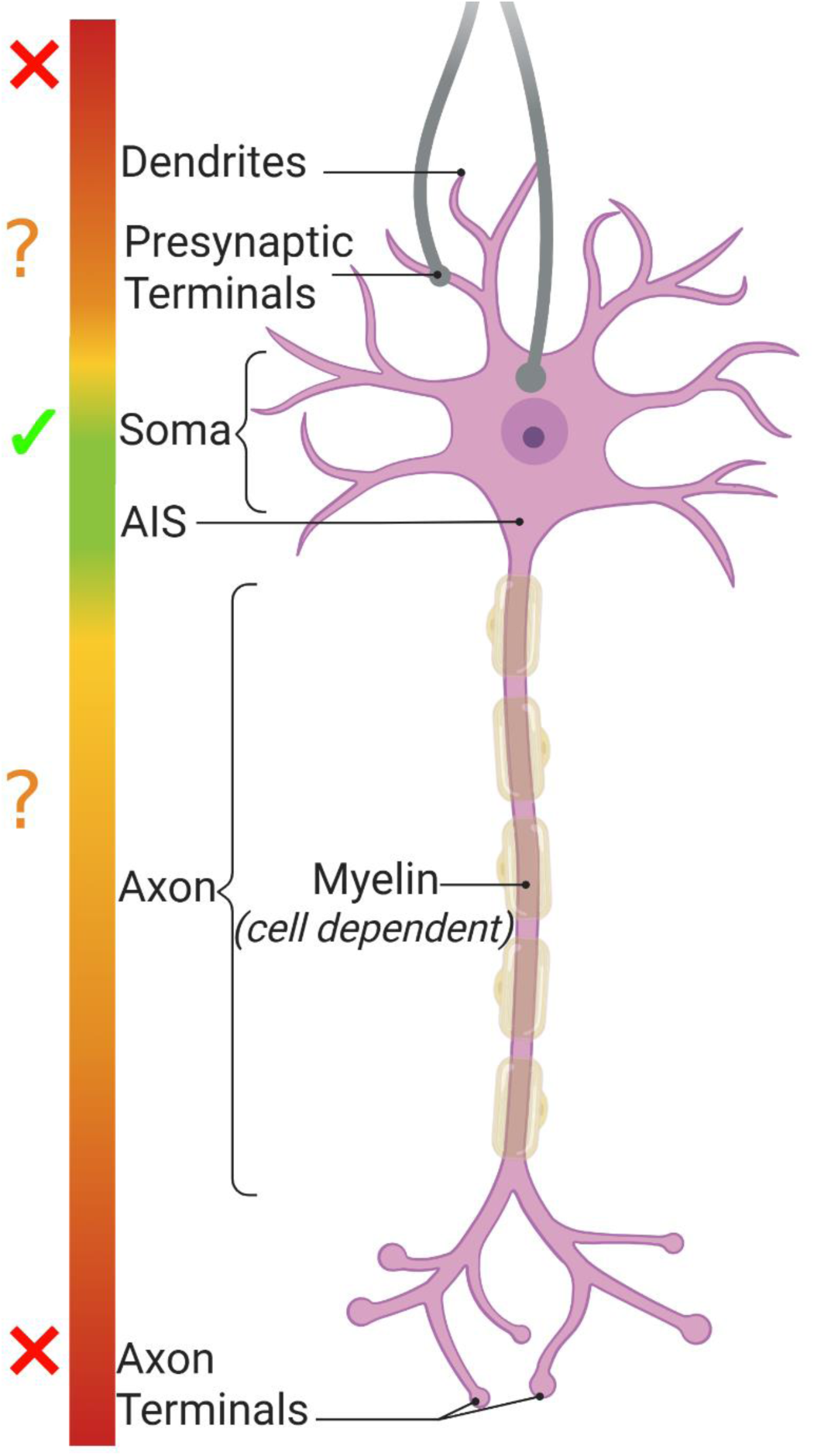
Spatial organization of neuronal compartments relevant to patch-SCP. Illustration of somatic, axonal, and synaptic compartments that contribute to electrophysiological signals measured during patch-clamp recording. In patch-SCP, proteomic recovery is expected to be enriched for somatic (green) and some proximal material (orange), whereas proteins localized to distal axons, presynaptic terminals, or specialized membrane domains may be under-represented (red) despite functional electrophysiological evidence of their activity. This spatial organization highlights the importance of compartment-aware interpretation when linking electrophysiology to single-neuron proteomic measurements. The schematic is not intended to represent the precise anatomy of any specific neuronal subtype.

Our proposed framework maps patch-clamp recording outcomes to distinct interpretations, defining what electrophysiological context can and cannot be meaningfully associated with the recovered proteome. Electrophysiology provides “true-positive” measurements of ion channel and synaptic function, whereas proteomic measurements are conditioned by retrieval efficiency; our framework provides a principled basis for interpreting agreement and disagreement between the two.

### Gigaseal Preservation: A Quantitative Bridge Between Proteomics and Electrophysiology

To test our framework, we sought to maintain the gigaseal during neuron relocation and to determine if recordings measured under this condition could predict proteome recovery (i.e. protein identifications). Figure 3A shows a representative image of neuron #4 and its current-clamp responses in the brain slice, taken prior to the initiation of the retrieval process. *In situ* recordings can provide valuable neurophysiological information (e.g. cell-typing by AP firing patterns) but have not been utilized in patch-SCP to quantify or qualify neuron retrieval (e.g., soma size and integrity or viability once withdrawn). Figure 3B shows that careful relocation of the soma to the slice surface did not disrupt the whole-cell configuration (n = 3), as evidenced by stable hyperpolarizing steps. The passive membrane properties for each neuron were initially assessed *in situ* (i.e. in the acute brain slice) and then reassessed during retrieval near the slice surface (Figure 3C). Our results show that while physiological changes can occur during retrieval, the gigaseal can be maintained well enough to reassess passive properties, such as resting membrane potential (V_M_; a proxy for health), capacitance (C, proportional to soma size), and resistance (R_M_; associated with membrane integrity). Importantly, total protein identifications correlated with the log-transformed capacitance (F = 1577, p < 0.05, adjusted R² = 0.998, n = 3, Y = 884.2*X + 197.7), providing a means of linking soma size to proteome recovery (Figure 3D). In contrast, log-transformed R_M_ showed no significant correlation with protein identifications (F = 0.748, p > 0.05; Figure 3E). These results indicate that soma size, which is proportional to capacitance, plays a more direct role in protein recovery than R_M_, which primarily reflects the ability of ions to move across the surface of the neuronal membrane (and is inversely proportional to size). Notably, over 1500 proteins were identified between these neurons, and at least 1700 total proteins were detected in each (Figure 3F). This consistent ‘core’ proteome included membrane-associated proteins (*APP*, *PRNP*, *APOE*), ion channel subunits (*SCN2A*, *CACNA2D1*, *GABRA1*, *GRIN1*), and GPCRs (*GRM3*, *GRM5*, *CHRM1*) (Figure 3G; Table S1).

**Figure 3.**
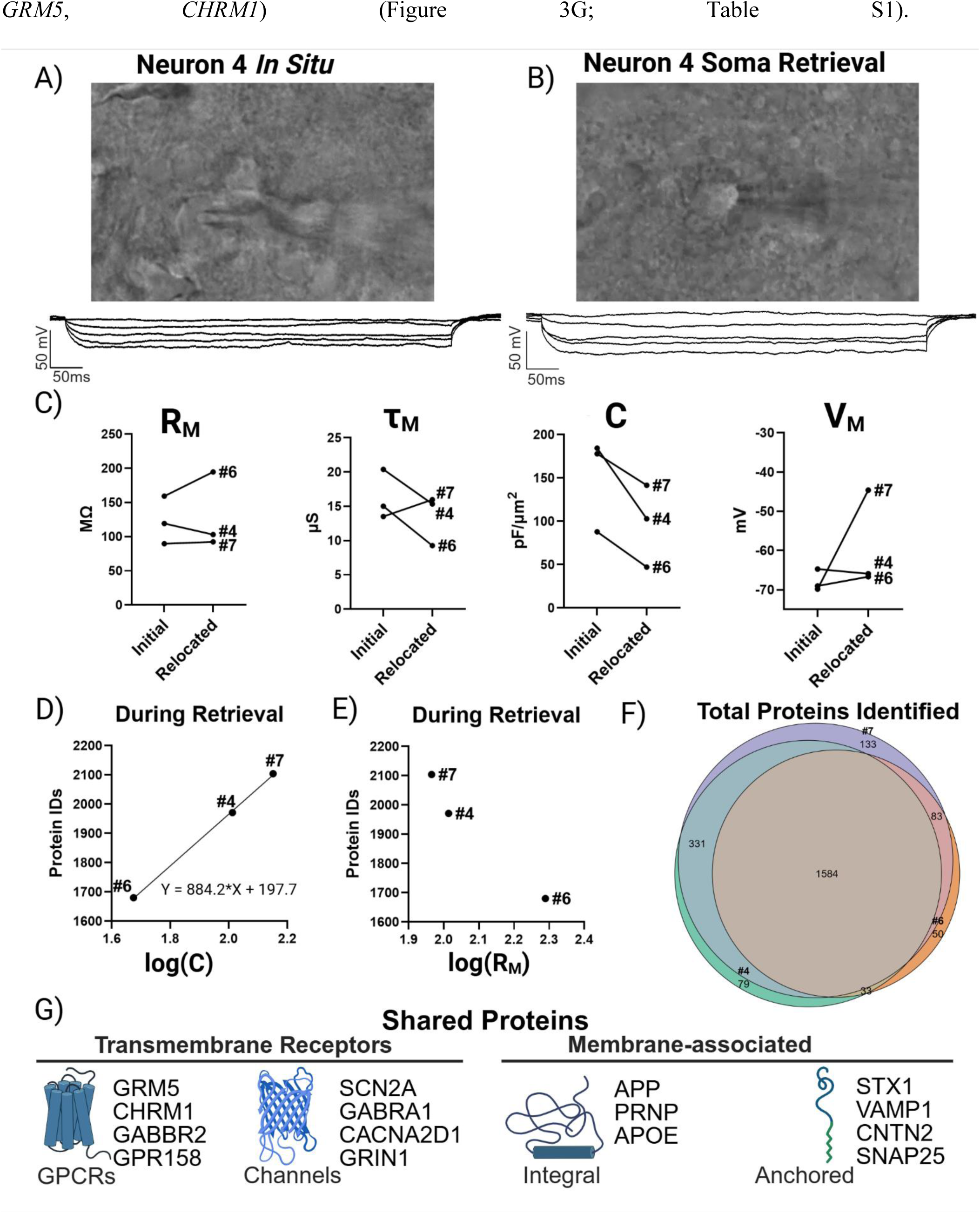
Gigaseal preservation links neuron size to proteome recovery. (A) Differential interference contrast (DIC) image and current clamp recording of neuron #4 before retrieval (i.e. *in situ*). (B) Soma of neuron #4 relocated to the slice surface while maintaining whole-cell access. (C) Ladder plots depicting a change in selected passive membrane properties from initial recordings and during retrieval of each neuron (n = 3). Passive properties were determined for each neuron from linear fits and extrapolated at I = 0 pA using NeuroExpress[15]. (D) Protein identifications correlated with log-transformed C, linking soma size to proteome yield. (E) No correlation was observed with log-transformed membrane resistance (R_M_). (F) Venn diagram showing a core proteome (>1500 proteins) shared across gigaseal-preserved neurons. (G) Examples of consistently recovered proteins, including *APP*, *PRNP*, *APOE*, *SCN2A*, *GRIN1*, and *GRM3*. Abbreviations: R_M_, membrane resistance; V_M_, resting membrane potential; τ_M_, membrane time constant; C, capacitance.

Given the association between a passive property like the capacitance of the soma being retrieved and proteome size, we next asked whether the collective integrity of the active membrane properties can be used to infer the specific neuronal functions of recovered proteins and more generally, biological insight. We reasoned that if the overall robustness of each soma’s action potential indicates the degree of physical disruption incurred, it might also reflect the probability that the retrieval would be enriched for proteins underlying synaptic signaling and neuronal identity. In support of this hypothesis, we found that relocating neurons to the surface of the slice did not prevent depolarizing currents from inducing action potentials. Indeed, we observed a range of effects on neuronal spiking in response to the retrieval processes (Figure 4A), demonstrating that the active membrane properties could generally be preserved during retrieval as to elicit a response from the soma. Representative videos of soma retrieval for each neuron are provided as Supporting Information (Videos S1–3), showing live differential interference contrast (DIC) imaging during withdrawal. Neuron #4 represents an ideal retrieval- a large, stable soma with no major issues (Video S1) in which depolarizing steps reliably induced consistent action potentials. By comparison, retrievals of neurons #6 and #7 were more challenging and both displayed altered active properties that are likely correlated with membrane disruption or damage near the axon hillock. Neuron #7 gently entered the electrode (Video S2) but still generated several action potentials with reduced amplitude during depolarizing steps, which is consistent with leak currents and a compromised membrane. In contrast, the partial aspiration of the contents of neuron #6 (Video S3) in the whole-cell configuration produced a more drastic outcome, with current-clamp recordings showing only a single spike during depolarization.

**Figure 4.**
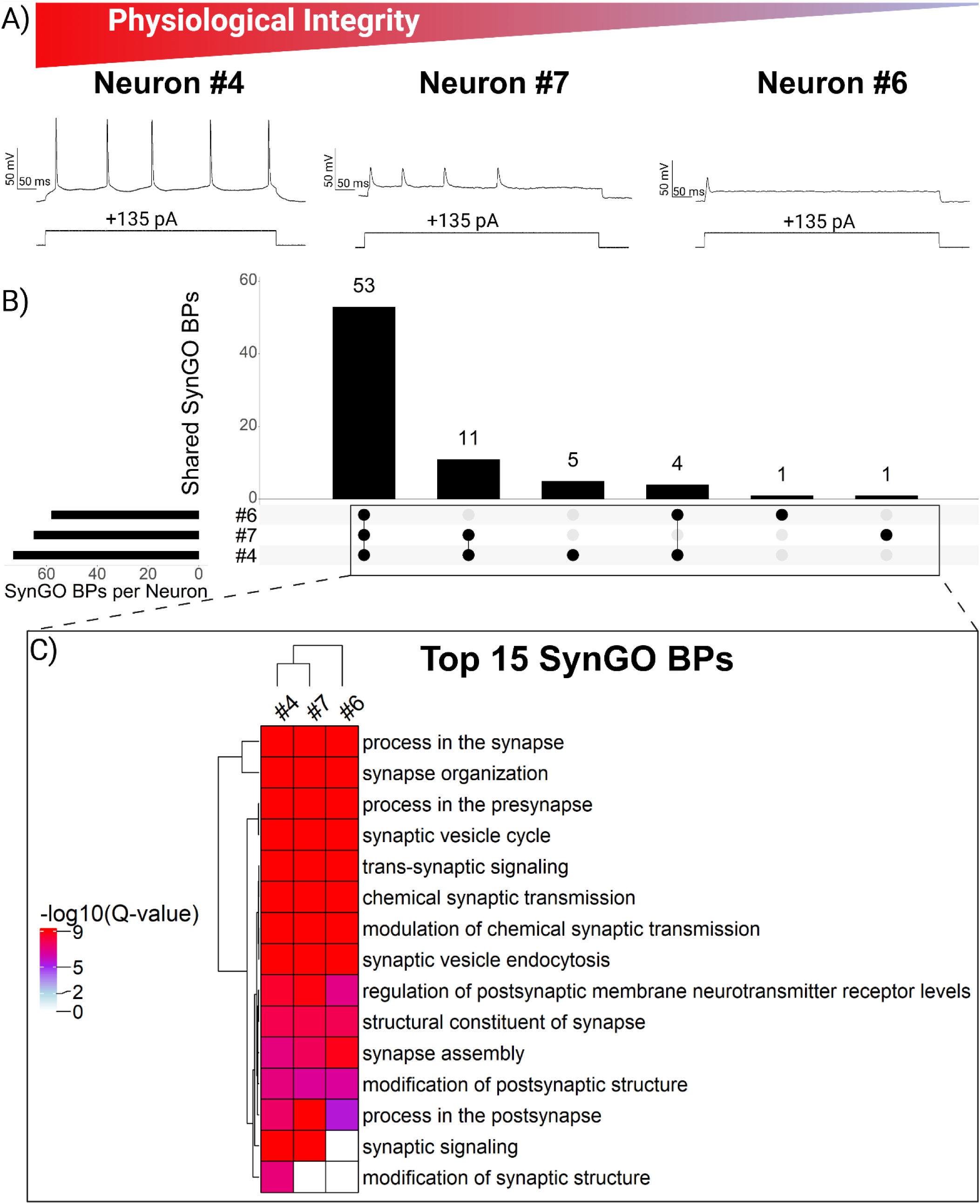
Preservation of active properties during retrieval is associated with recovery of synaptic proteins. (A) Action potential firing profiles illustrating differences in retrieval integrity. Neuron #4 retained stable spiking, neuron #7 showed reduced amplitude (consistent with current leak), and neuron #6 produced only a single spike following partial aspiration. (B) UpSet plot quantifying overlap of synaptic biological process (BP) terms (annotated by SynGO) across each neuron. (C) Heatmap of top BP terms, sorted by enrichment significance. Color intensity reflects statistical significance (-log_10_ adjusted Q-value; Q-value < 0.05 cutoff). Neuron #4 exhibited the greatest diversity of enriched terms, while neuron #6 lacked significant enrichment for synaptic signaling, despite many protein identifications.

To evaluate the impact of the soma retrieval process (as reflected by spike integrity) on the recovery of proteins with distinct neuronal functions, we utilized SynGO, a curated database tailored for analyzing gene ontologies of compartments and biological processes specific to synapses[16]. SynGO analysis revealed 53 biological processes (BPs; Figure 4B; Table S2) and 24 cellular components (CCs; Table S3) significantly enriched (Q-value < 0.05) across the proteomes of neurons retrieved under a gigaseal. Consistent with our finding that neuron size influences protein recovery, the smallest neuron (#6) exhibited the fewest enriched cellular component terms (27, versus 32 in #4 and 29 in #7). A similar trend was observed for biological processes (BPs), where the best-retrieved large neuron (#4) showed the greatest diversity of GO terms. Neurons #4 and #7 shared many features, including overlapping protein identifications (Figure 3F), detection of somatostatin (Table S1), enrichment of BPs such as synaptic signaling and organization (Figure 4C), and recovery of proteins from specialized CCs such as the active zone and anchored synaptic membrane components (Table S3). However, despite being the largest neuron by both electrophysiological and proteomic measurements, GO analysis of neuron #7 produced the fewest unique BP terms. Interestingly, the neuron with the most compromised active properties(#6) lacked significant enrichment of proteins associated with synaptic signaling (Q-value > 0.05), despite detection of several ion channel subunits such as *SCN2A* and *GABRA1* which are known to be involved in neurotransmission.

These results suggest that the electrophysiological properties that reflect neuron size (i.e. capacitance) can guide interpretation of proteomic recovery and challenge the common assumption that higher protein counts necessarily reflect deeper biological insight. Instead, our data indicate that the physiological condition of soma retrieval (reflected in the integrity of neuronal spiking during relocation), may be associated with the recovery of synaptic proteins. Preserving access to the soma during patch-SCP retrieval could provide future studies with an approach to either validate proteomic recovery across workflows or enhance the interpretability of appropriately powered functional investigations. For properties like current density, which normalizes the total ion current to the size of a neuron, accurate correlations with a specific proteomic profile (e.g. the relative abundance of ion channels, GPCRs, or transporters) will likely require measurements during soma retrieval.

### Retrieval Loss Decouples Proteomic Measurements from Electrophysiology Recordings

Given the complex nature of the patch-clamp technique, we maximized the efficiency of our small-scale gigaseal preservation study by indiscriminately retrieving all patch-SCP attempts for analysis. This “shotgun” approach to collection allowed us to ask whether the electrophysiological measurements made prior to neuron retrieval from the slice also provide insight into protein recovery. We categorized these retrievals as soma that 1) did not form a gigaseal, 2) lost the gigaseal during collection, or 3) were visibly and completely torn/aspirated during collection.

Figure 5A summarizes the protein identifications for each patch-SCP outcome, which can be classified by the failure to establish the whole-cell patch-clamp configuration (red) or by when the gigaseal was last intact, before (orange) or during soma retrieval (green). We chose to include neurons that were visually confirmed to be torn or aspirated into the electrode during retrieval (grey; Figure 5B) as negative controls. Despite having been characterized by electrophysiological recordings in the brain slice, these compromised samples produced the fewest detected proteins of all categories. Analysis of passive membrane properties measured prior to retrieval from the brain slice showed that neither log-transformed capacitance nor R_M_ correlated with protein identifications (p > 0.05; n = 6; Figures 5C-D). These results suggest that while recordings performed *in situ* provide valuable physiological insight, mechanical retrieval can introduce sample loss that decouples proteomic measurements from electrophysiological context, limiting how directly they can be co-interpreted. By comparison, retrievals without a gigaseal (i.e. soma isolated by micropipette without recordings) also suffered from variability but could yield similarly large proteomes, with 1,400-2,300 proteins detected per soma. More broadly, these results illustrate that retrieval of the soma from the brain slice can be so disruptive that initial electrophysiology recordings no longer reflect the collected material. We estimate that 25-50% of the soma could be lost over the course of collection (Figure 3C), although additional studies with a larger sample size are required to determine the extent of retrieval loss.

**Figure 5.**
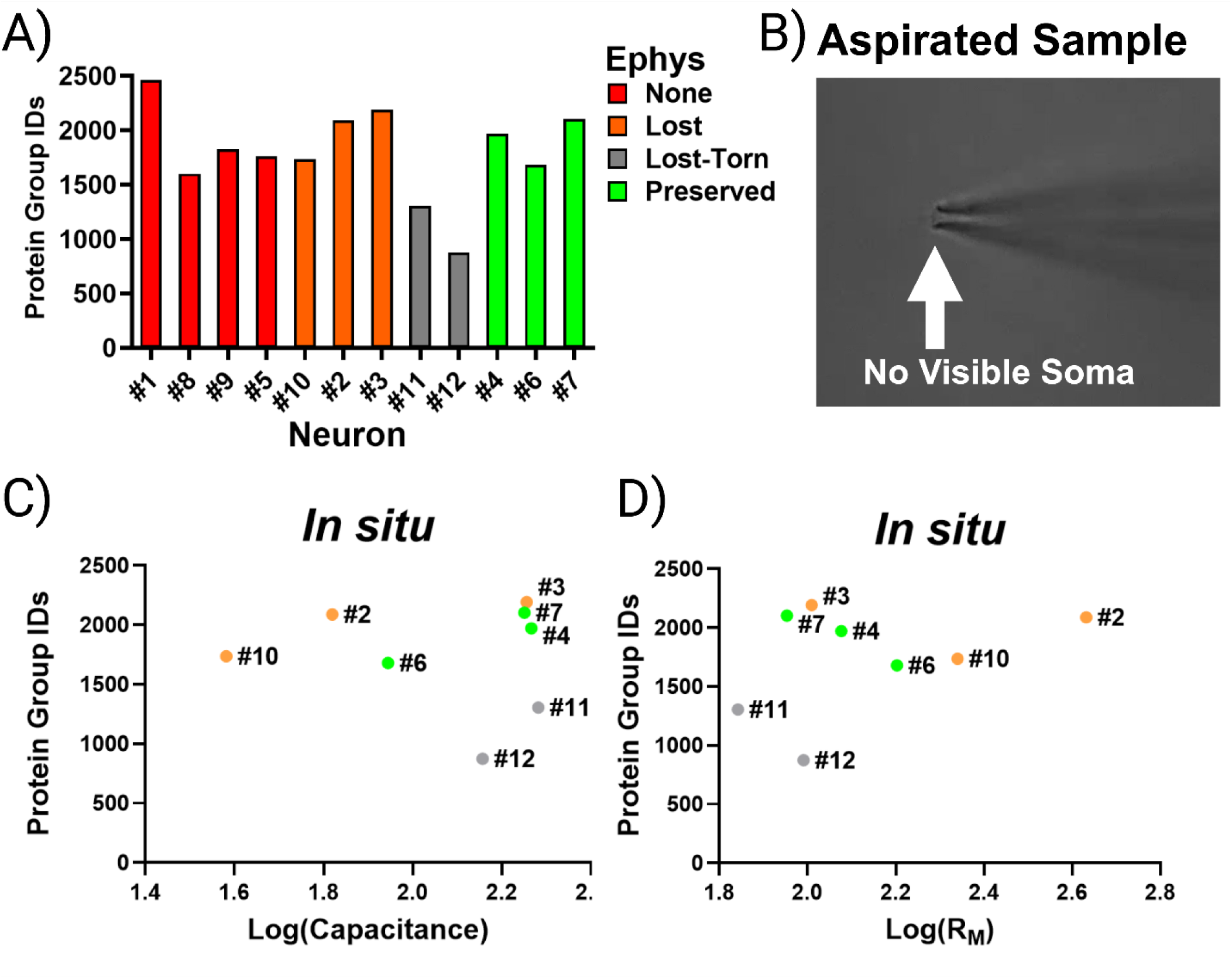
Proteome recovery is not associated with recordings performed *in situ*. (A) Protein identifications across all neurons, grouped by patch outcome. (B) Example of a “torn” neuron that was aspirated during retrieval, yielding few proteins. (C-D) Log transformations of capacitance and R_M_ measured prior to retrieval did not correlate with protein identifications (p > 0.05, n = 6). Colors denote the success-level of electrophysiological characterization throughout the patch-SCP process.

These data provide several key insights as a preliminary assessment of our SCP-MS capabilities. First, we saw that robust *in situ* recordings cannot compensate for severe retrieval loss, which can only be assessed by preserving the gigaseal during soma retrieval. We also observed that *in situ* recordings are not a prerequisite for achieving a large number of protein identifications, suggesting that our methods are sensitive enough to recover thousands of proteins from any neuron that remains visibly intact at the tip of the recording pipette. Lastly, torn samples still showed impressive protein recovery, demonstrating that as patch-SCP methods have become more sensitive, protein identifications have become less reliable indicators of sample quality.

### Comprehensive Neuron Retrieval: A Strategy For Evaluating Patch-SCP Variation

Recognizing that retrieval loss has the potential to uncouple SCP from patch-clamp recordings, we wanted to determine if our patch-SCP method could detect evidence of a soma that has been compromised. Indeed, principal component analysis (PCA) of all neurons (Figure 6A) revealed clear separation of extreme cases. Torn neurons (#11 and #12, grey) clustered apart from all others, consistent with their categorization as a poor retrieval (demonstrated in Figure 5B). Neuron #6, retrieved under a gigaseal but with compromised active properties, grouped more closely with neurons lacking gigaseals (#5, #8, #9; red), suggesting that its proteome resembled a coarse retrieval- large in terms of protein identifications but lacking key qualitative features, such as enrichment for synaptic signaling. Conversely, neuron #1 (no gigaseal) more closely resembled neurons with successful initial brain slice recordings (#2 and #3; orange). These results suggest that neuronal structures can be lost or retained in ways that are almost imperceptible when performing patch-SCP.

**Figure 6.**
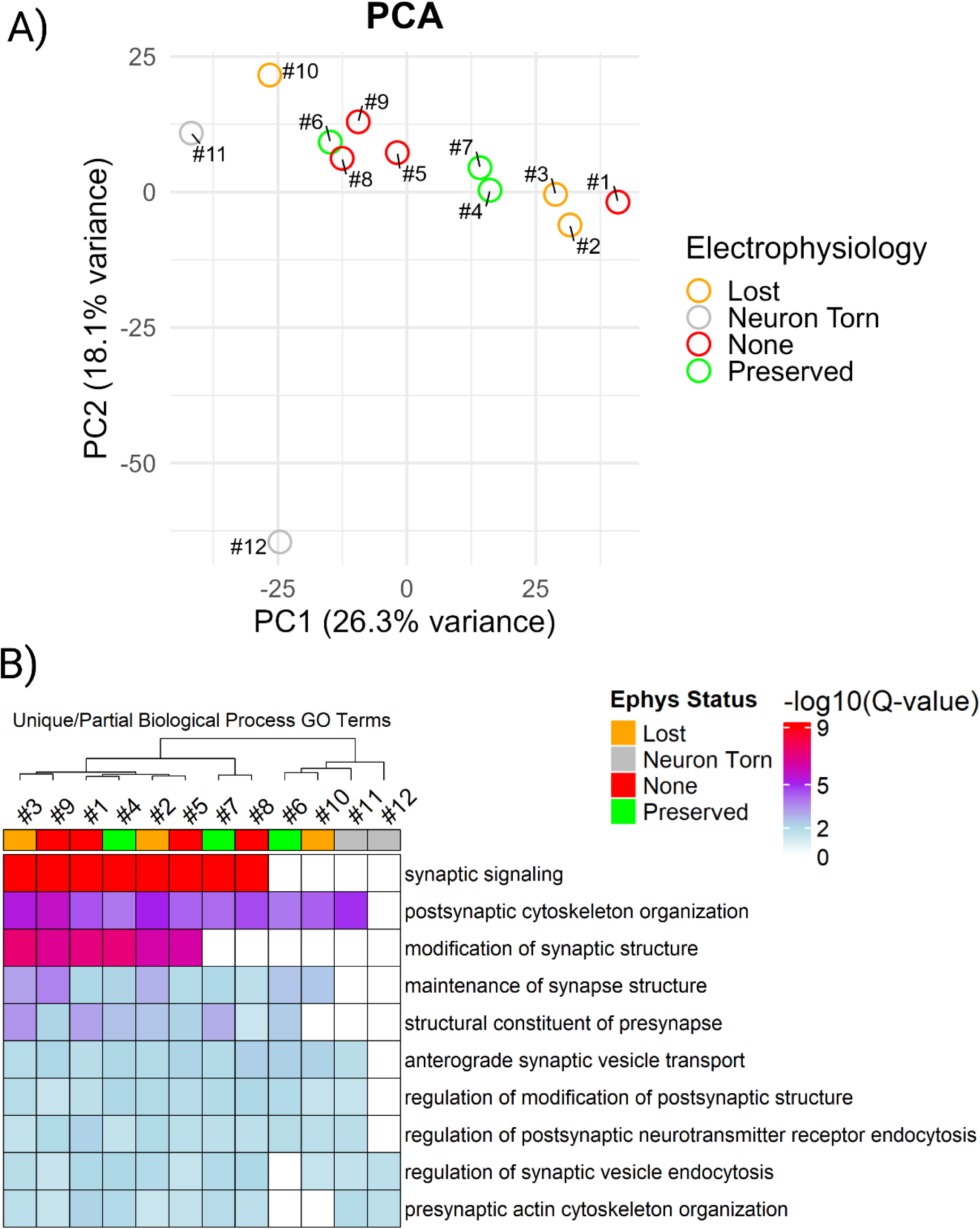
Comprehensive analysis can distinguish between high- and low-context retrievals. (A) PCA of single-neuron proteomes revealed clustering by factors other than patch outcome. Torn neurons (grey) separated from others, while neuron #6 (gigaseal but poor spiking) and neuron #10 (gigaseal lost during retrieval) clustered with cells that failed to form gigaseals (red). (B) Heatmap of SynGO-enriched biological processes which shows that synaptic enrichment varied across neurons, independent of outcome category.

Because proteomic profiles varied substantially within and across patch-clamp outcome categories, our data demonstrate that categorical patch-clamp outcome alone does not reliably predict the biological content of single-neuron proteomes. To determine whether the variation in proteomic components observed across retrieval outcomes reflected differences in biological insight, we performed GO analysis on the proteome of each neuron using SynGO[16]. Across all neurons, we identified a total of 23 cellular components and 45 biological processes that were consistently enriched (Figure S2; Q-value < 0.05). Notably, SynGO CC enrichments clustered along retrieval category (Figure S3), similar to the pattern observed in the PCA plot, rather than by the number of identified proteins. Detection of a core set of synaptic processes suggests that consistent proteomic coverage by our patch-SCP workflow is enabled by the convergence of several complementary factors: (i) meticulous neuron retrieval, (ii) sample preparation methods that minimize loss, and (iii) the sensitivity of high-performance data independent acquisition (DIA) mass spectrometry. Our results also demonstrated the benefit of a proteome-centric approach for exploratory patch-clamp studies, particularly when the decision to develop or abandon SCP methods hinges on the efficient generation of small, actionable data sets. Studies designed to answer a targeted biological question could use this approach to determine sample inclusion criteria. For example, analysis of partially shared biological processes revealed that several neurons(#6 and #10) lacked significant enrichment for synaptic signaling despite being characterized by electrophysiology (Figure 6B). By clustering with torn neurons(#11 and #12), which serve as internal standards for compromised retrieval, these results demonstrate that variability during retrieval should not be assumed to be mitigated by the success of *in situ* electrophysiology recordings or the detection of a specific number of proteins. Instead, the recovery of synaptic information for each sample should be assessed and compared against the relative sampling depth of the patch-SCP workflow.

In summary, developing new SCP methods can require the rapid analysis of many samples that have been optimized across multiple variables, such as isolation technique, sample preparation, and MS acquisition. While rigorous, this approach is not compatible with manual patch-clamp electrophysiology, which is why we pursued the more practical strategy of proteome-centric shotgun analysis of all retrieval attempts. Here, the unpredictable nature of the protein losses incurred during soma retrieval can be partially revealed by a comprehensive analysis of all patch outcomes, where comparisons across categories can resolve compromised retrievals in a way that protein identification cutoffs cannot. We emphasize that characterization of neurons by SCP alone is not a substitute for characterization by patch-SCP. Rather, the analysis of all samples makes it possible to (i) benchmark retrieval (given soma are visible) and sample-handling performance against proteomic readouts, and (ii) supplement molecular information that can corroborate hypotheses (e.g., pathway enrichment or receptor detection) when functional data are limited.

### Detecting Transmembrane Proteins with Patch-SCP

Ion channels, GPCRs, and transporters represent the most exciting targets for patch-SCP because they play a direct role in mediating neurophysiology. Therefore, we wanted to assess whether our patch-SCP workflow was robust enough to detect low abundance and hydrophobic transmembrane proteins.

We evaluated the detection of membrane proteins in each sample by creating ion channel, GPCR, and transporter recovery lists, using a combination of SynGO annotations and curated gene tables from IUPHAR-DB, a peer-reviewed resource of pharmacologically relevant protein families [17, 18]. Figure 7 illustrates that our workflow detected diverse classes of ligand- and voltage-gated ion channels, including GABA_A_ receptor subunits (e.g., *GABRA1*, *GABRB2*, *GABRG2*), AMPA and NMDA receptor subunits (*GRIA1-3*, *GRIN1*, *GRIN2B*), and multiple components of Ca_v_2.2 channels (*CACNA1B*, *CACNA2D1-3*, *CACNB4*, *CACNG8*). We were able to identify several ion channel subunits with distinct functional roles across samples, regardless of patch-clamp retrieval category. In all neurons, we detected the α subunit of the ligand-gated GABA_A_ ion channel, which produces inhibitory postsynaptic currents (IPSCs), as well as *CACNB4* the auxiliary β subunit of the voltage-gated Ca_v_2.2 channels, which regulate excitability and neurotransmitter release[19]. The voltage-dependent anion channel *VDAC1* was also detected in all neurons, although this protein is associated with the maintenance of a potential gradient across the outer mitochondrial membrane and, unlike ion channels found on the plasma membrane, is not expected to be found in low-abundance[20]. *SCN2A*, the alpha subunit of Na_v_1.2 that initiates AP propagation [21], was detected in all neurons except #11, which was torn during retrieval. Its near-universal detection is consistent with its expected expression within the axonal initial segment and across CNS neurons[22]. We also observed a peptide that maps to transient receptor potential channels, though ambiguous assignments precluded its confident inclusion in Figure 7.

**Figure 7.**
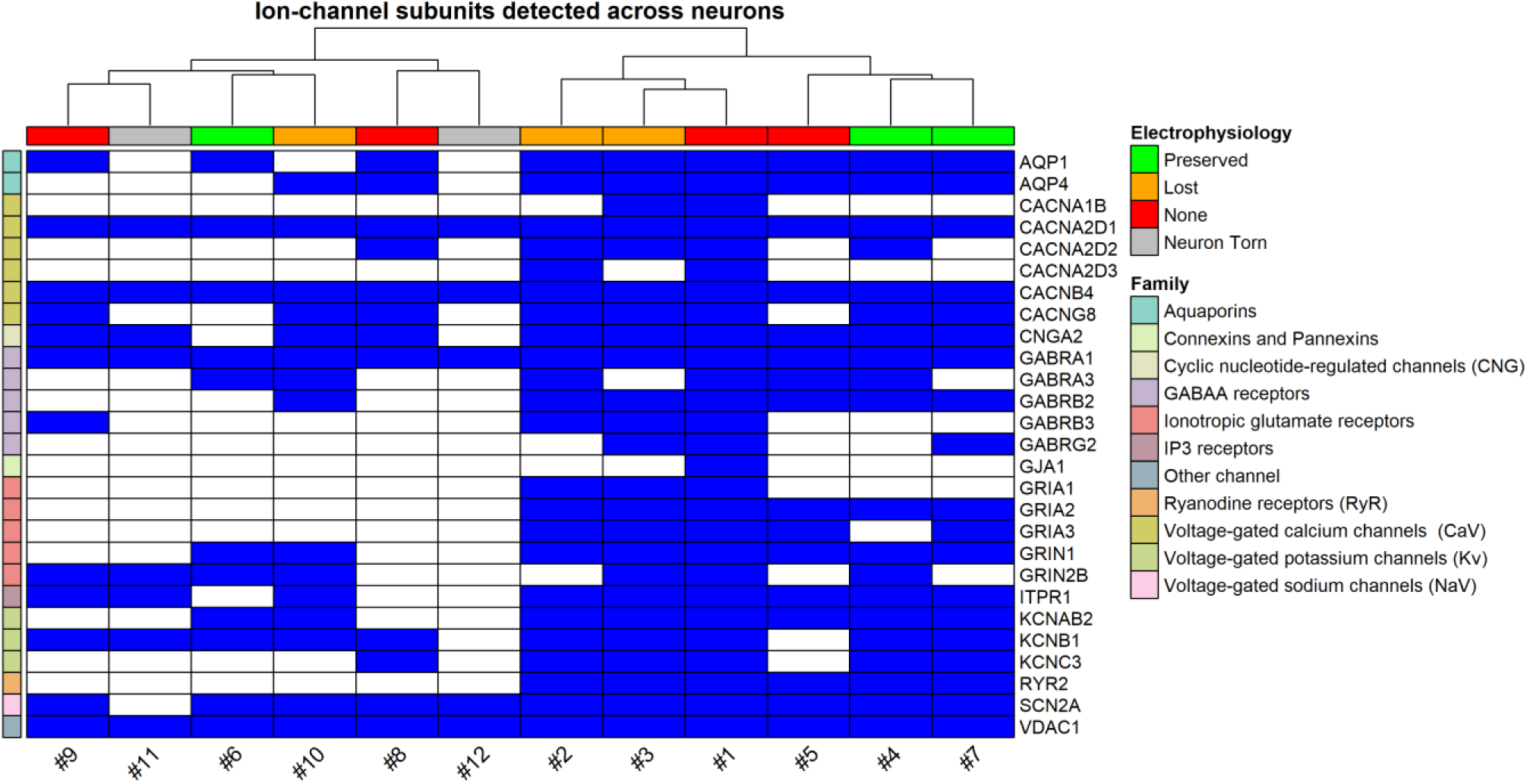
Recovery of ion channels across neurons retrieved by patch-SCP. Binary heatmap showing detected ion channel subunits (blue) across individual neurons. Rows are grouped by channel family (e.g. sodium, calcium, glutamate, GABA). Columns are annotated by the success-level of electrophysiological characterization during the patch-SCP process. Neurons with gigaseals generally retained broader sets of ion channel subunits, although more subunits were recovered some gigaseal-lost neurons than in preserved ones. Torn neurons consistently showed the poorest recovery.

Overall, ion channel, GPCR (Figure S4), and transporter (Figure S5) detection patterns seemed to vary with electrophysiological characterization. The higher-context retrievals shown in Figure S3, which often included neurons with initial electrophysiological recordings, also tended to retain a broader diversity of transmembrane proteins, whereas the lower-context cluster of torn or severely compromised soma consistently showed reduced recovery. Importantly, ion channel, GPCR, and transporter recovery did not always scale with electrophysiological success at the time of retrieval; in some cases, more transmembrane proteins could be found among neurons that lost the whole-cell configuration during retrieval than neurons with continuous whole-cell access. This could reflect variability in the recovery of membrane compartments, as ion channels are both low in abundance and spatially restricted, although our sample size is insufficient to determine whether these differences are technical or biological.

### Limitations and Future Directions

The mechanics of soma retrieval presents an underexplored limitation when interpreting patch-SCP in acute brain slices. As highlighted by others, variability in the proteins identified by mass spectrometry can undermine statistical analyses and weaken the impact of robust electrophysiological findings[7, 8]. Accordingly, our goal was to explore how soma retrieval outcomes influence patch-SCP analysis. To achieve this, we developed a patch-SCP framework that included indiscriminate soma collection, which allowed us to interpret our data along two orthogonal dimensions: proteome yield (a quantitative axis) and biological integrity of recovered compartments (a qualitative axis).

We hypothesized that maintaining whole-cell access (an indicator of careful and minimally disruptive extraction) during soma retrieval would preserve cellular material and thereby improve the depth and biological fidelity of the recovered proteome. We found that measurements of somal capacitance made during retrieval were associated with protein identifications, suggesting that the intrinsic properties of the soma could reflect a “quantitative axis” (in this case, the amount of neuronal material being collected; Figure 3D). However, soma retrieval introduces a qualitative axis in which damage or loss of specific compartments can selectively distort the biological processes detected, even when protein identifications remain high. We report that maintaining access to the soma can make the integrity of samples easier to identify during retrieval, but due to the three dimensional organization of acute brain slices, variability cannot be completely mitigated. A general example of stochastic retrieval variability involves the isolation of synaptic terminals where pre- and postsynaptic elements are physically coupled by structural proteins. While synaptic structures can be manually retrieved with the soma, it is typically not possible to isolate only the local terminal of an individual cell, and thus most retrievals of local termini contain some synaptic proteins from the presynaptic cell they are connected with. Cortical neurons also exhibit substantial heterogeneity in axonal morphology and specialization, including differences in branching patterns and myelination status. Because patch-SCP isolates only the soma and immediately adherent material (Figure 2), the present approach cannot resolve these structural differences at the level of individual retrieved neurons.

While our framework proved useful for exploring our patch-SCP capabilities, it also revealed areas that need improvement to enhance interpretability. The recovery of ion channel subunits could vary widely, ranging from 4 proteins in the case of a torn neuron (#12) to as many as 27 proteins in a neuron retrieved without a gigaseal (#1). This is a promising start for a method, but future refinements to the workflow are necessary to increase reliability and reproducibility, particularly to correlate specific neuronal functions with proteomic profiles. For example, finding an association between channel kinetics and the relative abundance of ion channels, GPCRs, or transporters would likely require stringent inclusion criteria (i.e. gigaseal preservation and enrichment of specific CC/BPs). To this end, we detected subunits of several major ion channel complexes (e.g., *SCN2A*, *CACNA2D1–3*, *GABRA1*) and neuropeptides such as somatostatin (SST) and VGF, but we did not consistently recover complete channel assemblies from individual somas and failed to detect several GPCR families of high biological interest (e.g., opioid, adrenergic, serotonergic, and CRF_1_ receptors). Because many ion channels, receptors, transporters, and neuropeptides are synthesized in the soma but are rapidly trafficked to specialized membrane domains, incomplete recovery may reflect either technical limitations of soma retrieval or biological compartmentalization beyond the cell body[23]. In cases where a protein’s function is primarily localized to distal axonal or synaptic compartments, soma-focused sampling is therefore expected to bias detection toward underrepresentation. Because long-range neuronal processes are severed during slice preparation, the absence of distal proteins cannot be attributed to either slice truncation or mechanical loss during soma retrieval. Resolving this distinction will require benchmarking across larger datasets.

In this study, we chose to focus on soma retrieval. However, patch-SCP is a versatile technique that was originally used to sample cytoplasmic content. In principle, rupturing the membrane patch while maintaining the gigaseal means that, over the course of a patch-clamp recording, the contents of the cytoplasm and the recording electrode dialyze one another. This process may be exploited to make cytoplasmic patch-SCP more efficient, where the only requirement for analysis may be successfully going whole-cell (see Figure S1) for a certain amount of time, which is already standard for pharmacological investigations. Such cytoplasmic sampling may allow neurons to be characterized by the expression of higher abundance, synapse-specific protein markers or intracellular signaling molecules like protein kinase C, although the exact target will depend on the region of interest and the experimental conditions. Notably, previous cytoplasmic and somal patch-SCP analyses have relied on data-dependent acquisition (DDA) whereas our mass spectrometry analysis was performed using data-independent acquisition (DIA). Although whole-proteome DIA provides a strong starting point for exploratory patch-SCP, the consistent detection of low-abundance, hydrophobic, or spatially restricted proteins will likely require additional optimization. To enhance sensitivity and reproducibility for predefined molecular targets, future studies may benefit from improved sample preparation strategies and the integration of targeted or hybrid acquisition modes, such as DIA workflows augmented with parallel reaction monitoring (PRM).

It is important to consider how SCP depth is influenced by the same anatomical constraints that make acute slice electrophysiology possible. Preparation of 300 µm coronal slices necessarily severs long-range projections prior to patching, constraining retrieval to the soma and proximal compartments. Beyond this global truncation, the local environment may also impact soma recovery. Neurons embedded within dense synaptic or dendritic networks could experience greater mechanical resistance to withdrawal, although retrieval would still be possible if these neurons were located close to the slice surface where tissue integrity is most disrupted. In the present dataset, we visibly recovered 10 of 12 pyramidal somas and did not observe systematic differences in extraction success based on intrinsic electrophysiological properties. However, due to our limited sample size we cannot conclude whether retrieval efficiency varies across defined neuronal subtypes or morphologies. For example, neuron #1 yielded an unusually large proteome and was located near the slice surface (Video S4), suggesting that local geometry may influence recovery in ways not fully captured by electrophysiological metrics alone. Resolving whether such cases represent biological variability or technical bias will require larger, subtype-defined datasets and orthogonal markers of neuronal identity (e.g., reporter lines, *post hoc* labeling, or morphological reconstruction). Within the scope of this proof-of-concept study, our framework is therefore best suited for benchmarking retrieval integrity and molecular coverage in relatively large neurons, rather than for making definitive claims about subtype-specific extraction efficiency.

A final patch-SCP limitation concerns the spatial fidelity of electrophysiological measurements themselves. In neurons with extended morphology, somatic voltage clamp cannot uniformly control membrane potential across distal dendrites or the axon initial segment due to electrotonic separation, access resistance, and non-uniform channel distributions[24]. As a result, currents recorded at the soma often reflect spatially filtered contributions from compartments that are not well clamped. This limitation is particularly relevant for patch-SCP, because many ion channels and receptors that shape neuronal excitability are enriched in distal or specialized domains that may neither be electrically well controlled nor physically recovered with the soma. Consequently, even when electrophysiological recordings demonstrate channel activity, neither the magnitude nor the kinetics of those currents can be assumed to map quantitatively onto somatic protein recovery. This constraint does not diminish the utility of patch-SCP, but it shows the importance of aligning the biological question, the electrophysiological measurement, and the compartment sampled for proteomic analysis. Consistent with this challenge, recent Patch-seq work using 955 neurons from the adult mouse motor cortex demonstrated that hybrid statistical–biophysical modeling can use gene-expression profiles to predict parameters of conductance-based (Hodgkin–Huxley) models fit to electrophysiological recordings[25]. However, cell-type diversity and model mismatch continued to complicate efforts to link molecular measurements to distributed channel mechanisms, even when multimodal data and many samples are available.

## Conclusions

We present an interpretive framework for integrating electrophysiological and proteomic measurements at the single-neuron level in acute brain slices for patch-SCP. By monitoring neuronal properties during retrieval, we demonstrated that parameters such as capacitance and spike integrity provide practical indicators of proteome recovery. Our results show that retrieval mechanics, rather than *in situ* electrophysiology alone, limit whether proteins associated with excitability and synaptic function are recovered in the single-cell proteome. Recognizing this limitation motivated a comprehensive inclusion strategy in which all patch outcomes were analyzed, providing retrieval context, benchmarking sample quality, and reducing the risk of overinterpreting protein counts in isolation. At the same time, the incomplete recovery of certain membrane-associated proteins highlights the need for continued optimization of retrieval and sample preparation workflows. Together, these findings establish shotgun patch-SCP as a proof-of-concept approach and define a practical framework for interpreting how electrophysiological context and retrieval variability shape single-neuron proteomic measurements. By clarifying these limitations, this work offers guidance for the design and interpretation of future patch-SCP studies in semi-intact brain circuits, particularly those aimed at relating molecular composition to neuronal identity and signaling under defined experimental conditions.

## Methods and Materials

### Electrophysiology

All animal studies conformed to NIH Guidelines for the care and use of laboratory animals and were approved by the Scripps Research Institutional Animal Care and Use Committee (IACUC; protocol no. 09-0006). Wistar rats (75 days of age, Charles River) were given access to food and water *ad libitum*. Acute brain slices and electrophysiological recordings were performed as previously described [10, 26–31]. Briefly, rats were anesthetized with isoflurane before cervical dislocation and surgical brain isolation. Coronal mPFC slices (300 μm) were prepared in an ice-cold high-sucrose cutting solution (sucrose 206 mM; KCl 2.5 mM; CaCl_2_ 0.5 mM; MgCl_2_ 7 mM; NaH_2_PO_4_ 1.2 mM; NaHCO_3_ 26 mM; glucose 5 mM and HEPES 5 mM) using a Leica VT1200S vibratome. Slices were incubated in 95% O_2_/5% CO_2_ equilibrated artificial cerebrospinal fluid (aCSF; NaCl 130 mM; KCl 3.5 mM; NaH_2_PO_4_ 1.25 mM; MgSO_4_·7H_2_O 1.5 mM; CaCl_2_ 2.0 mM; NaHCO_3_ 24 mM and glucose 10 mM), first at 37°C for 30 minutes, then at room temperature for another 30 minutes. Slices were then transferred to a recording chamber (Warner Instruments) and superfused at a flow rate of 2-4 ml/min. Whole-cell patch-clamp recordings were performed in neurons present in the medial subdivision of the PrL and clamped at −70 mV using a Multiclamp 700B amplifier, Digidata 1440A and pClamp10. Patch pipettes (3-6 MΩ) were filled with an internal solution composed of the following (in mM): 145 KGluconate, 0.5 EGTA, 2 MgCl_2_, 10 HEPES, 2 Mg-ATP and 0.2 Na-GTP. The intrinsic membrane properties and excitability of L2/3 neurons were determined in aCSF in current-clamp mode using a step protocol comprised of 500 ms hyperpolarizing and depolarizing steps in 5 or 10 pA increments (see [32]). NeuroExpress software (version 19.4.09.) developed and provided by A. Szücs was used for analysis[15]. During withdrawal of the patched neuron, light negative pressure (-50 to -140 mmHg) was applied through the pipette and adjusted to facilitate soma retrieval while preserving the high-resistance seal. This gentle suction helped maintain gigaseal integrity as the soma was relocated toward the slice surface. Following retrieval, an image of the pipette tip was taken above the slice to confirm whether an intact soma was present before proceeding to sample processing. All available videos of the patch-SCP attempts described in this study have been made publicly available in their full form on Zenodo (DOI: 10.5281/zenodo.18189812).

### Sample Processing and LC-MS/MS Acquisition

Micropipettes containing the retrieved neurons were immediately transferred to a 384-well non-binding microplate (Greiner) containing 15 µL of 0.02% n-dodecyl-β-D-maltoside (DDM) in UHPLC-grade water (ThermoFisher). To release the soma and its contents into the surfactant solution for subsequent processing, the pipette tip was carefully snapped in the well plate, which was stored on dry ice throughout the collection process. Samples were digested at 37°C for 2 hours using 7 ng of sequencing-grade Trypsin (Promega), quenched with 2 µL of 0.1% formic acid, and stored at -20°C for later processing. Peptide separation was performed using a Vanquish Neo UHPLC system (ThermoFisher) coupled to a 25 cm × 75 μm IonOpticks Aurora XS column with an integrated emitter. Samples were loaded directly and separated using a 36-minute gradient at 400 nL/min. A large wash cycle was included between runs to minimize carryover. MS analysis was performed on an Orbitrap Astral mass spectrometer (ThermoFisher Scientific) operating in data-independent acquisition (DIA) mode with FAIMS Pro using a single compensation voltage (CV = -50). Survey scans were acquired at 240,000 resolution (at m/z 200) over a mass range of 400-1000 m/z. DIA windows were 20 m/z wide with overlapping edges. MS1 AGC target was set to 800%, with a maximum injection time of 50 ms. Fragmentation was performed using higher-energy collisional dissociation (HCD) with normalized collision energy (NCE) of 25.

### Bioinformatic and Statistical Analysis

Raw DIA files were analyzed using DIA-NN v1.8.1 in library-free mode with the “match-between-runs” option enabled[33]. Searches were performed against the UniProt Mus musculus reference proteome (downloaded 2024), with oxidation selected as a variable modification. Up to two missed tryptic cleavages were allowed. Decoys were generated using the default reversed-sequence approach. Protein identifications were filtered at 1% false discovery rate (FDR) at both the precursor and protein group level. For all downstream analysis, only protein groups that passed both filters (i.e., those appearing in DIA-NN’s report.pg_matrix.tsv) were retained. Proteomic quantification was performed using DIA-NN’s MaxLFQ implementation, which integrates peptide-level signals into protein-level intensity estimates across samples. Enrichment terms for cellular component (CC) and biological process (BP) categories were derived from SynGO [16], a manually curated synaptic GO ontology database. Gene set enrichment analyses were performed on gene lists derived from DIA-NN’s protein-level output. For SynGO analysis, proteins were annotated based on gene symbols and GSEA filtering was performed under stringent conditions. The threshold for statistical significance for SynGO analyses was Q-value of < 0.05. Per-cell gene lists were extracted from the DIA-NN output file report.pg_matrix.tsv using detected/not detected as thresholds after 1% protein-level FDR filtering. GO term enrichment and clustering analyses were conducted using R 4.3.1 and visualized with packages including ComplexHeatmap, ggplot2, and UpSetR. Custom scripts for figure generation and SynGO-based analysis are available at https://github.com/LarryThePharmacologist. Ion channel, GPCR, and transporter annotation lists were generated using curated gene families from SynGO [16] and IUPHAR-DB [17, 18]. The raw mass spectrometry data and search files have been deposited to the ProteomeXchange Consortium (http://proteomecentral.proteomexchange.org) via the MassIVE partner repository with the MassIVE dataset identifier MSV000099156 and ProteomeXchange dataset identifier PXD068359.

## Supporting information

Tables containing protein-level DIA-NN v1.8.1 and SynGO output

## Author Information

### Author Contributions

Conceptualization: JY, JD, LR, MR; Methodology: LR, AM, LS, RV, BT, SK, CB; Software Validation: SK, BT, CB; Formal Analysis: LR, LS, AM, BT, CB, SK; Investigation: LR, BT, SK, CB; Resources: JY, MR, CB; Data Curation: LR, SK, CB; Writing-Original Draft: LR, LS, JY, MR, RV; Visualization: LR, JD, JY, MR; Supervision: AM, MR, RV, JY; Project Administration: AM, MR, JY; Funding Acquisition: MR, JY.

## Supporting Information

The Supporting Information is available free of charge at https://pubs.acs.org/doi/xxxxxxxxxxx/.

Figure legends; schematic depicting the whole-cell patch-clamp process; UpSet plots of significant synaptic gene ontologies (SynGO); heatmap showing GPCR detection across samples; video legends (PDF)

Tables containing protein-level DIA-NN v1.8.1 and SynGO output. (XLSX)

Video S1. Gigaseal-preserved soma retrieval of neuron #4 (AVI).

Video S2. Partial preservation of soma integrity for neuron #7 (AVI).

Video S3. Partial aspiration of neuron #6 during soma retrieval (AVI).

Video S4. Soma retrieval of neuron #1 without electrophysiological characterization (AVI).

## Acknowledgements

We thank Claire Delahunty for feedback on the manuscript; Titus Jung for assistance with DIA-NN analysis and data processing; Patrick Garrett for developing tools that supported data exploration; Casimir Bamberger, Attila Szücs, and Michal Bajo for helpful discussions on mass spectrometry, proteomics, and electrophysiology; Myka Kairs for helpful suggestions on video processing; and Sabyasachi Baboo and Natalie Turner for technical guidance. We are also grateful to Antonio Pinto and the Scripps Research Institute Multi-omics Core Facility for their insight and support.

## Abbreviations

SCP: single-cell proteomics
patch-SCP: patch-clamp single-cell proteomics
DIA: data-independent acquisition
MS: mass spectrometry
MS/MS: tandem mass spectrometry
hiPSC: human induced pluripotent stem cell
mPFC: medial prefrontal cortex
GPCR: G protein-coupled receptor
Na_V_: voltage-gated sodium channel
Ca_V_: voltage-gated calcium channel
AMPA: α-amino-3-hydroxy-5-methyl-4-isoxazolepropionic acid receptor
NMDA: N-methyl-D-aspartate receptor
GABA: γ-aminobutyric acid
IPSC: inhibitory postsynaptic current
AP: action potential
V_M_: resting membrane potential
τ_M_: membrane time constant
C_cell_: cell capacitance
C_Soma_: soma capacitance
R_M_: membrane resistance
R_p_: pipette resistance
R_A_: access resistance
CC: cellular component
BP: biological process
GO: Gene Ontology
VDAC: voltage-dependent anion channel
SST: somatostatin
CRF_1_: corticotropin-releasing factor receptor 1
FAIMS: high-field asymmetric waveform ion mobility spectrometry.

## For Table of Contents Only

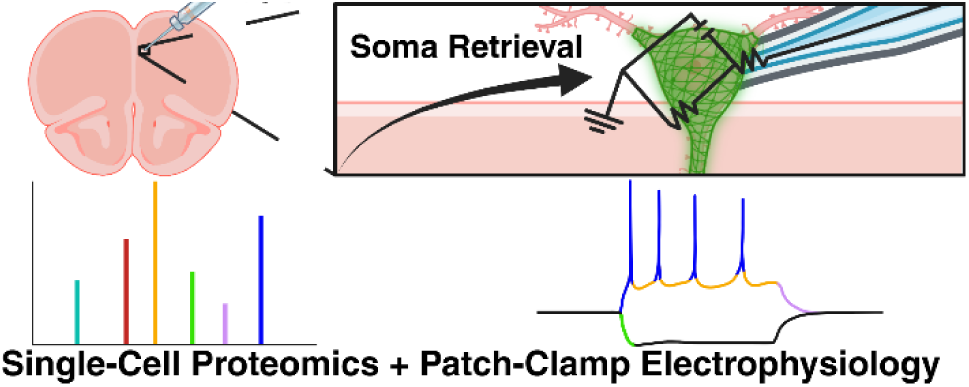

---

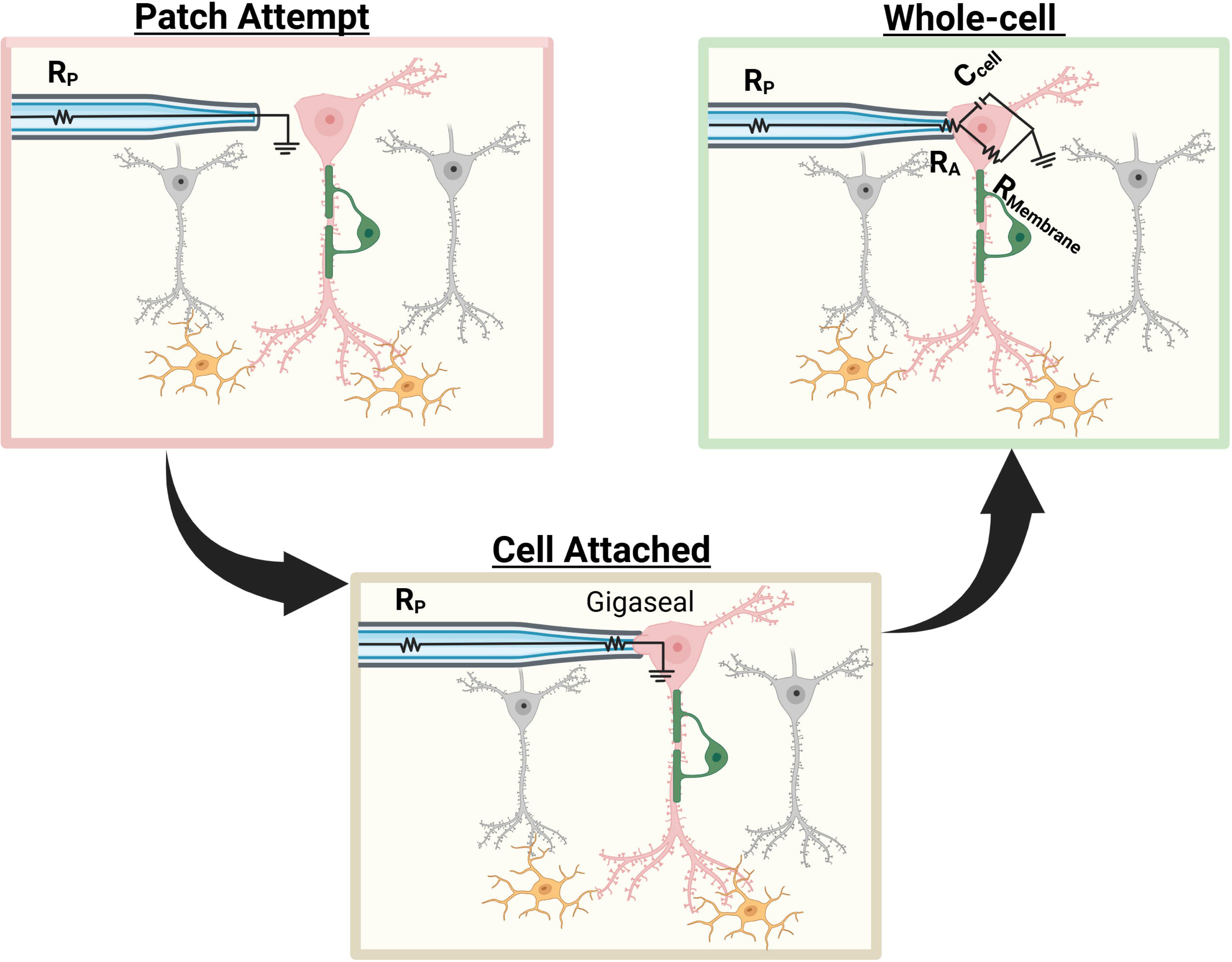

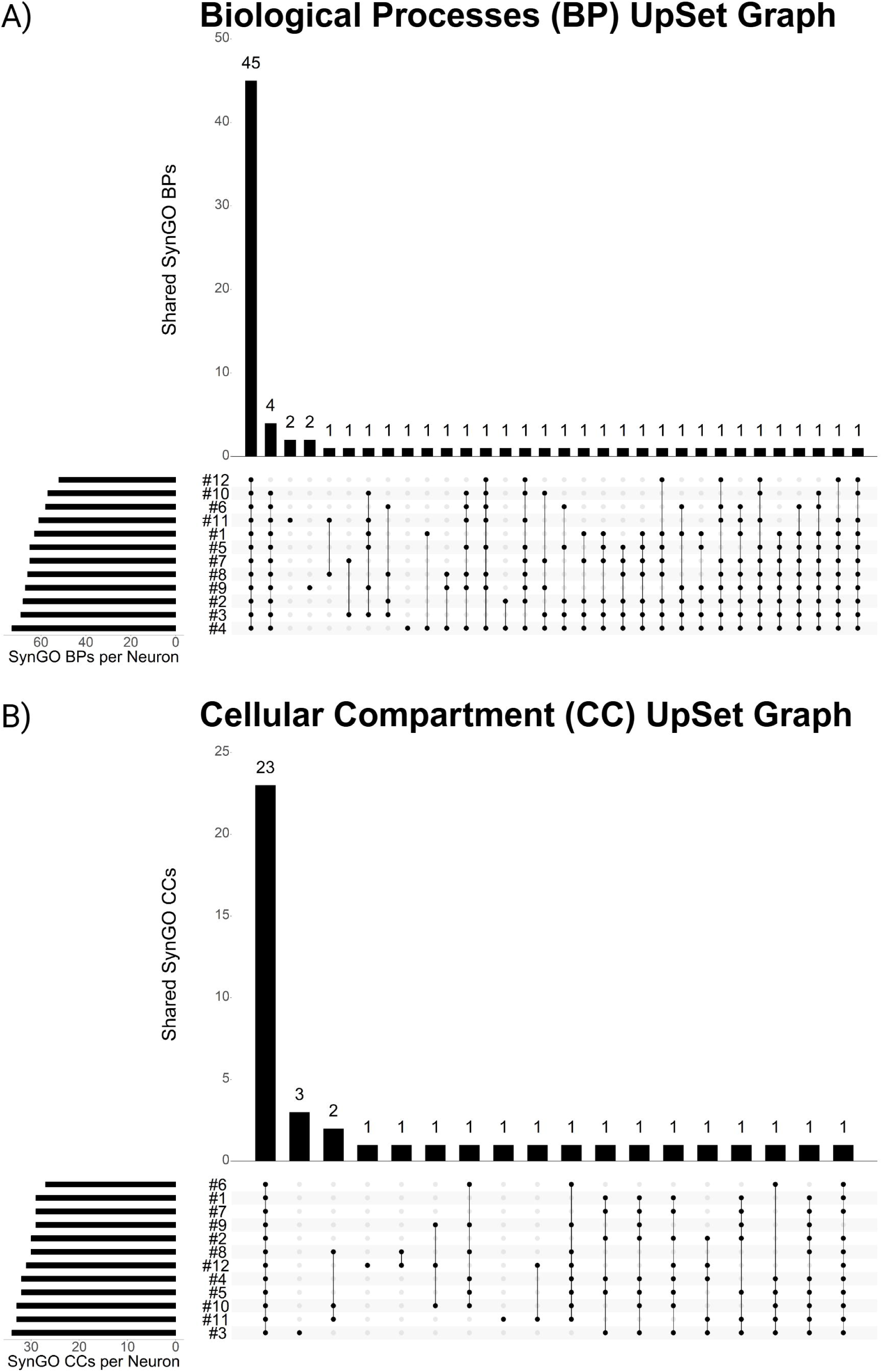

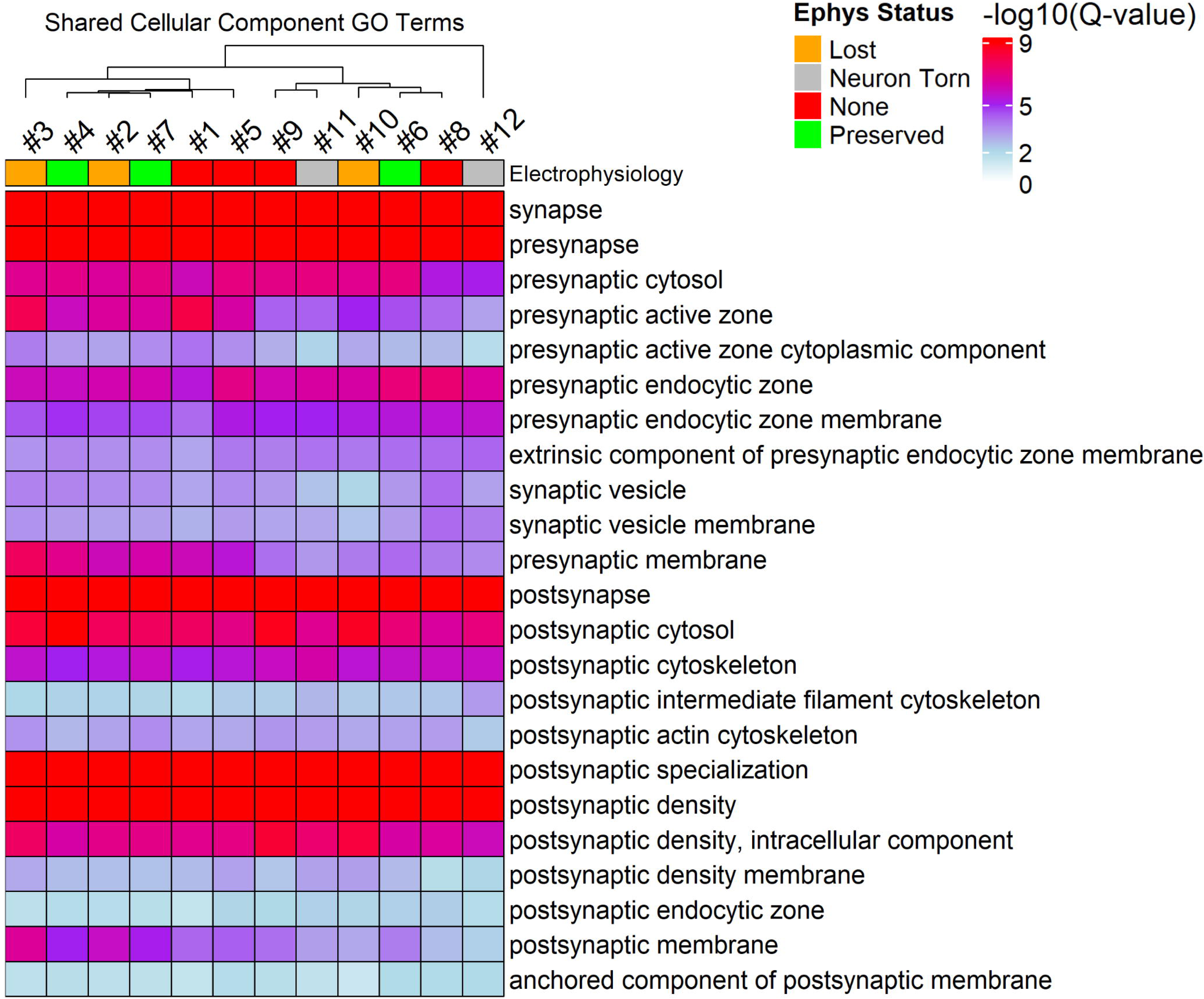

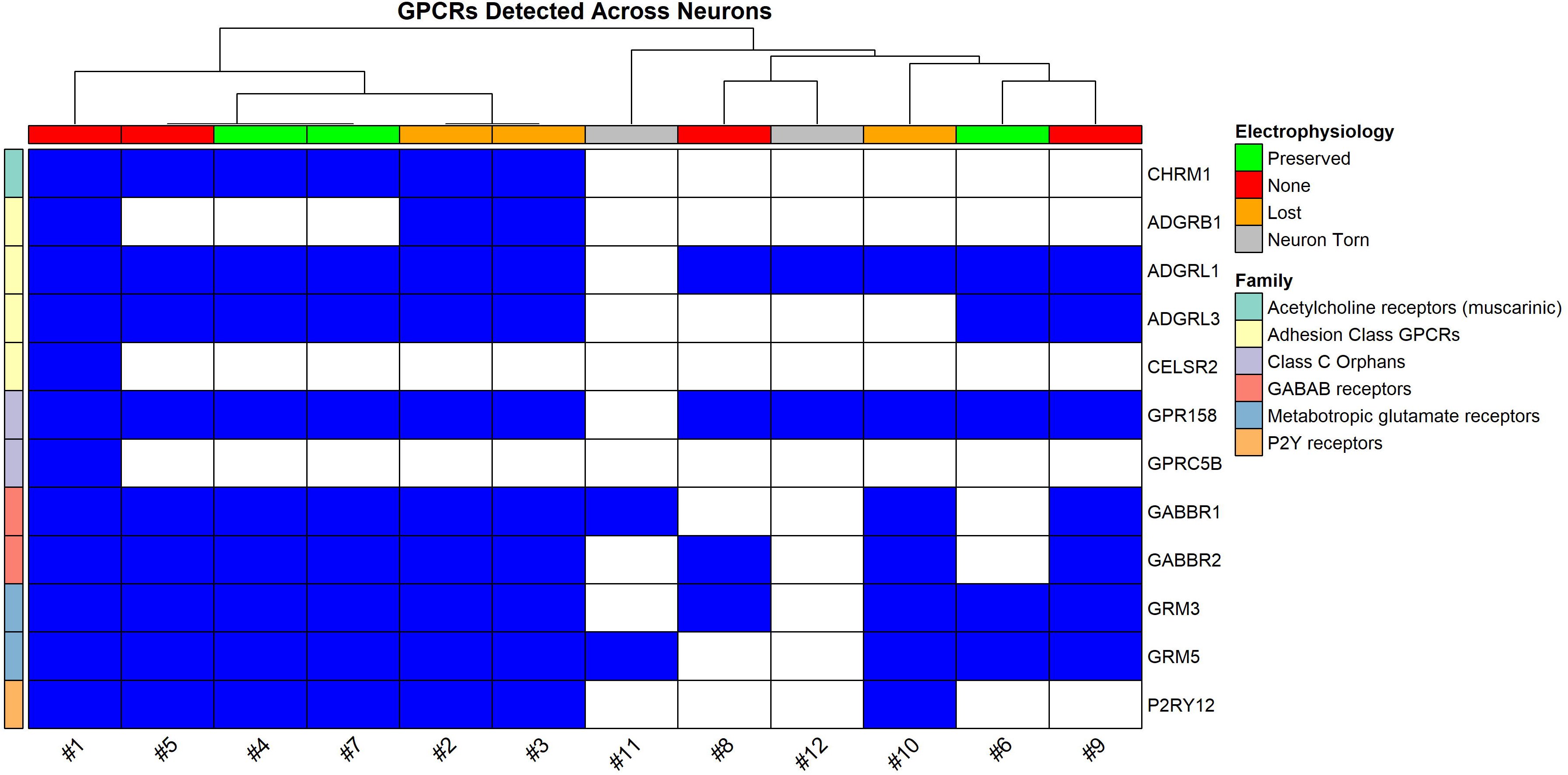

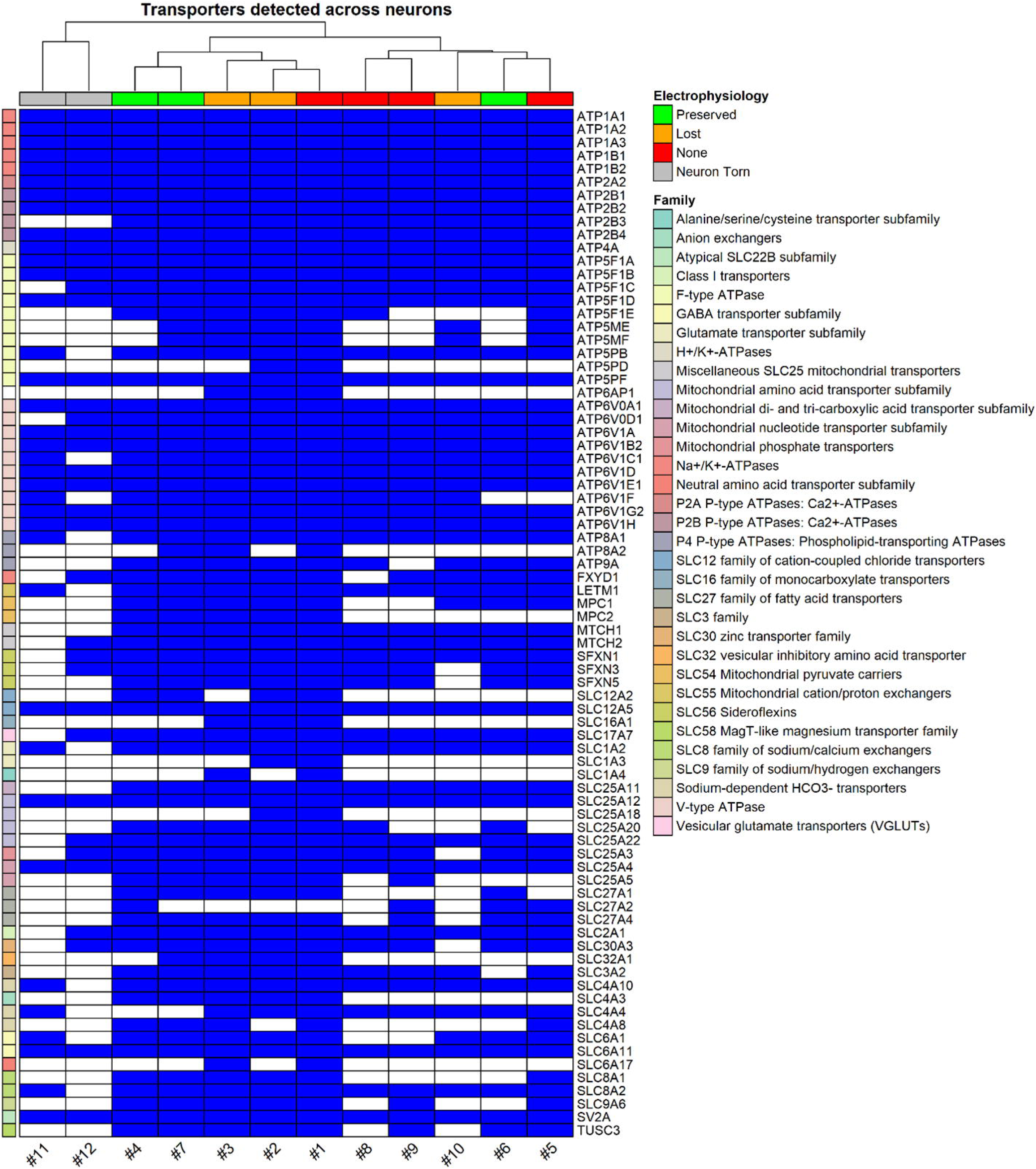

## References

1. Qiu, S., et al., Single-neuron RNA-Seq: technical feasibility and reproducibility. Frontiers in Genetics, 2012. 3.

2. Cadwell, C.R., et al., Electrophysiological, transcriptomic and morphologic profiling of single neurons using Patch-seq. Nat Biotechnol, 2016. 34(2): p. 199–203.

3. Hrvatin, S., et al., Single-cell analysis of experience-dependent transcriptomic states in the mouse visual cortex. Nat Neurosci, 2018. 21(1): p. 120–129.

4. Lipovsek, M., et al., Patch-seq: Past, Present, and Future. J Neurosci, 2021. 41(5): p. 937–946.

5. Lambolez, B., et al., AMPA receptor subunits expressed by single purkinje cells. Neuron, 1992. 9(2): p. 247–258.

6. Aerts, J.T., et al., Patch Clamp Electrophysiology and Capillary Electrophoresis–Mass Spectrometry Metabolomics for Single Cell Characterization. Analytical Chemistry, 2014. 86(6): p. 3203–3208.

7. Choi, S.B., A.M. Polter, and P. Nemes, Patch-Clamp Proteomics of Single Neurons in Tissue Using Electrophysiology and Subcellular Capillary Electrophoresis Mass Spectrometry. Anal Chem, 2022. 94(3): p. 1637–1644.

8. Lee, J., et al., Sex differences in single neuron function and proteomics profiles examined by patch-clamp and mass spectrometry in the locus coeruleus of the adult mouse. Acta Physiol (Oxf), 2024. 240(4): p. e14123.

9. Ghatak, S., et al., Single-Cell Patch-Clamp/Proteomics of Human Alzheimer’s Disease iPSC-Derived Excitatory Neurons Versus Isogenic Wild-Type Controls Suggests Novel Causation and Therapeutic Targets. Adv Sci (Weinh), 2024. 11(29): p. e2400545.

10. Patel, R.R., et al., Functional and morphological adaptation of medial prefrontal corticotropin releasing factor receptor 1-expressing neurons in male mice following chronic ethanol exposure. Neurobiol Stress, 2024. 31: p. 100657.

11. Daviu, N., et al., Neurobiological links between stress and anxiety. Neurobiology of Stress, 2019. 11: p. 100191.

12. Wu, C.C., et al., A method for the comprehensive proteomic analysis of membrane proteins. Nature Biotechnology, 2003. 21(5): p. 532–538.

13. Gatto, L., et al., Initial recommendations for performing, benchmarking and reporting single-cell proteomics experiments. Nature Methods, 2023. 20(3): p. 375–386.

14. Südhof, T.C., Towards an Understanding of Synapse Formation. Neuron, 2018. 100(2): p. 276–293.

15. Szücs, A., NeuroExpress program for analyzing patch-clamp data. ResearchGate, 2022.

16. Koopmans, F., et al., SynGO: An Evidence-Based, Expert-Curated Knowledge Base for the Synapse. Neuron, 2019. 103(2): p. 217–234.e4.

17. Alexander, S.P.H., et al., The Concise Guide to PHARMACOLOGY 2019/20: Ion channels. British Journal of Pharmacology, 2019. 176(S1): p. S142–S228.

18. Alexander, S.P.H., et al., The Concise Guide to PHARMACOLOGY 2023/24: G protein-coupled receptors British Journal of Pharmacology, 2023. 180(S2): p. S23–S144.

19. Zamponi, G.W., et al., The Physiology, Pathology, and Pharmacology of Voltage-Gated Calcium Channels and Their Future Therapeutic Potential. Pharmacological Reviews, 2015. 67(4): p. 821–870.

20. Shoshan-Barmatz, V., et al., Subcellular localization of VDAC in mitochondria and ER in the cerebellum. Biochimica et Biophysica Acta (BBA) - Bioenergetics, 2004. 1657(2): p. 105–114.

21. Hu, W., et al., Distinct contributions of Nav1.6 and Nav1.2 in action potential initiation and backpropagation. Nature Neuroscience, 2009. 12(8): p. 996–1002.

22. Garrido, J.J., et al., A Targeting Motif Involved in Sodium Channel Clustering at the Axonal Initial Segment. Science, 2003. 300(5628): p. 2091–2094.

23. Hille, B., Ion channels of excitable membranes. 3rd ed. 2001, Sunderland, Mass: Sinauer.

24. Armstrong, C.M. and W.F. Gilly, Access resistance and space clamp problems associated with whole-cell patch clamping, in Methods in Enzymology. 1992, Academic Press. p. 100–122.

25. Bernaerts, Y., et al., Combined statistical-biophysical modeling links ion channel genes to physiology of cortical neuron types. Patterns, 2025. 6(10).

26. Rodriguez, L., et al. Alcohol Dependence Induces CRF Sensitivity in Female Central Amygdala GABA Synapses. International Journal of Molecular Sciences, 2022. 23, DOI: 10.3390/ijms23147842.

27. Vlkolinsky, R., et al., Withdrawal from chronic alcohol impairs the serotonin-mediated modulation of GABAergic transmission in the infralimbic cortex in male rats. Neurobiol Dis, 2024. 199: p. 106590.

28. Varodayan, F.P., et al., Chronic ethanol induces a pro-inflammatory switch in interleukin-1β regulation of GABAergic signaling in the medial prefrontal cortex of male mice. Brain Behav Immun, 2023. 110: p. 125–139.

29. Athanason, A.C., et al., Chronic ethanol alters adrenergic receptor gene expression and produces cognitive deficits in male mice. Neurobiol Stress, 2023. 24: p. 100542.

30. Anjos-Santos, A., et al., Noradrenaline Modulates Central Amygdala GABA Transmission and Alcohol Drinking in Female Rats. Biol Psychiatry, 2025.

31. Guo, Y., et al., Scalable total synthesis of saxitoxin and related natural products. Nature, 2025. 646(8084): p. 351–357.

32. Patel, R.R., et al., Ethanol withdrawal-induced adaptations in prefrontal corticotropin releasing factor receptor 1-expressing neurons regulate anxiety and conditioned rewarding effects of ethanol. Mol Psychiatry, 2022. 27(8): p. 3441–3451.

33. Demichev, V., et al., DIA-NN: neural networks and interference correction enable deep proteome coverage in high throughput. Nature Methods, 2020. 17(1): p. 41–44.

