## Supporting Information for "Patch-Clamp Single-Cell Proteomics in Acute Brain Slices: A Framework for Recording, Retrieval, and Interpretation"

**Legends**

**Table S1 (XLSX).** Protein-level DIA-NN1.8.1 output (report.pg_matrix.tsv). Contains gene names, quantification values, and associated data for all analyzed neurons.

**Table S2 (XLSX).** SynGO Biological Process (BP) enrichment results for each neuron. Each column corresponds to a neuron, with rows representing enriched GO terms and statistical values. Stringent evidence filter selected.

**Table S3 (XLSX).** SynGO Cellular Component (CC) enrichment results for each neuron, structured as in Table S2.

**Video S1. Gigaseal-preserved soma retrieval of neuron #4 (AVI).** Live differential interference contrast (DIC) microscopy showing retrieval of neuron #4 from an acute rat mPFC brain slice while maintaining whole-cell access. The soma remains visibly intact at the pipette tip throughout relocation toward the slice surface. Electrophysiological recordings were performed before and during retrieval, as described in Figure 3 and Figure 4. Playback speed is increased (~7× real time) and the view is cropped to the microscope image for clarity. The full-length, real-time recording (including electrophysiology traces) is available on Zenodo (DOI: 10.5281/zenodo.18189812).

**Video S2. Partial preservation of soma integrity for neuron #7 (AVI).** Live DIC microscopy showing retrieval of neuron #7. The soma is collected with partial preservation of morphology but exhibits altered electrophysiological properties during relocation, consistent with membrane disruption near the axon initial segment.
Playback speed is increased (~7× real time); only the microscope view is shown. The complete recording is available on Zenodo (DOI: 10.5281/zenodo.18189812).

**Video S3. Partial aspiration of neuron #6 during soma retrieval (AVI).** Live DIC microscopy showing retrieval of neuron #6, in which partial aspiration of somatic contents occurred during withdrawal. This retrieval was associated with severely compromised active membrane properties and limited synaptic protein enrichment (Figures 4–6). Playback speed is increased (~7× real time); only the microscope view is shown. The complete recording is available on Zenodo (DOI: 10.5281/zenodo.18189812).

**Video S4. Soma retrieval of neuron #1 without electrophysiological characterization (AVI).** Live DIC microscopy showing retrieval of neuron #1, which was collected without successful formation of a gigaseal. Despite the absence of electrophysiological data, the soma was visibly intact at the pipette tip and yielded one of the largest proteomes in the dataset. Playback speed is increased (~7× real time); microscope view only. The full-length video is available on Zenodo (DOI: 10.5281/zenodo.18189812).

**Figure S1. Schematic of the patch-clamp process**. Diagram illustrating the transition from initial pipette contact ("Patch attempt", top left) to full whole-cell configuration (top right), which allows simultaneous electrical and cytosolic access to individual neurons in the brain slice. The circuit elements (pipette resistance R_P_, access resistance R_A_, membrane resistance R_Membrane_, and cell capacitance C_cell_) are shown only for the whole-cell configuration. The intermediate cell-attached configuration (gigaseal formed but membrane patch unruptured) is shown on the bottom. Created with BioRender.

**Figure S2.** **SynGO enrichment across single neurons.** (A) biological processes (BP) and (B) cellular component (CC) gene ontology (GO) terms across single-cell samples. A core set of 45 BP and 23 CC terms significantly enriched in all 12 neurons, indicating reproducible detection of synaptic proteins. GSEA Q-value significance cutoff < 0.05

**Figure S3.** **Shared SynGO CC enrichment across single neurons.** Heatmap of 20 SynGO cellular component (CC) gene ontology (GO) terms shared across single-cell samples. Clustering was performed by samples (columns) only. GSEA Q-value significance cutoff < 0.05

**Figure S4.** **GPCR detection across single neurons.** Binary heatmap showing presence (blue) or absence (white) of G protein-coupled receptors (GPCR) identified in individual neurons. Samples are annotated by the level of electrophysiological characterization (top bar). Green denotes whole-cell configuration was “Preserved”. Orange denotes whole-cell configuration was “Lost” during retrieval. Gray denotes whole-cell configuration was lost due to the neuron being “Torn” during retrieval. Red denotes no electrophysiological characterization due to a failure to form a gigaseal during the initial patch attempt. Clustering was performed by samples (columns) only.

**Figure S5.** **Transporter detection across single neurons.** Binary heatmap showing presence (blue) or absence (white) of protein transporters identified in individual neurons. Samples are annotated by the level of electrophysiological characterization (top bar). Green denotes whole-cell configuration was “Preserved”. Orange denotes whole-cell configuration was “Lost” during retrieval. Gray denotes whole-cell configuration was lost due to the neuron being “Torn” during retrieval. Red denotes no electrophysiological characterization due to a failure to form a gigaseal during the initial patch attempt. Clustering was performed by samples (columns) only.

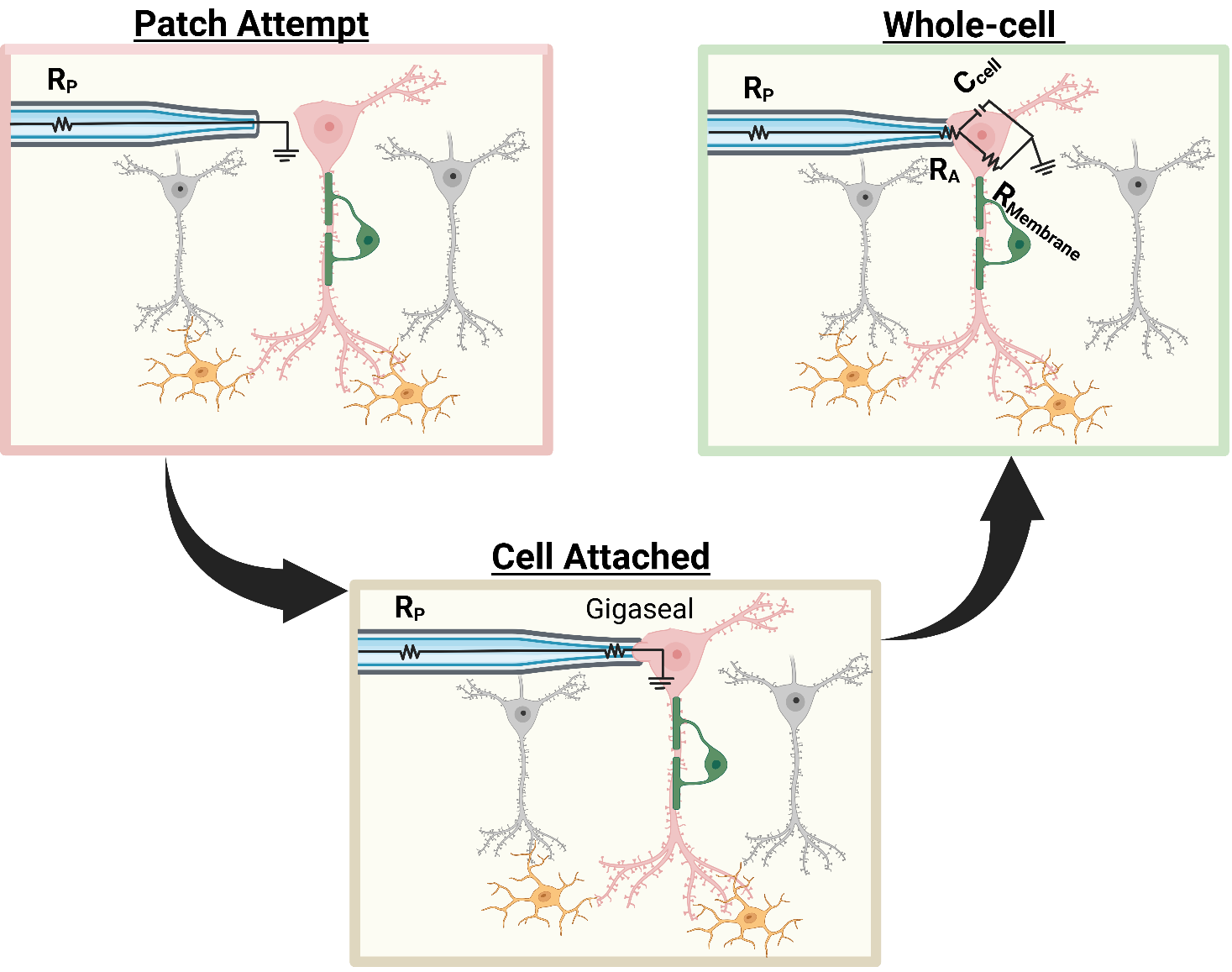

**Figure S1. Schematic of the patch-clamp process**. Diagram illustrating the transition from initial pipette contact ("Patch attempt", top left) to full whole-cell configuration (top right), which allows simultaneous electrical and cytosolic access to individual neurons in the brain slice. The circuit elements (pipette resistance R_P_, access resistance R_A_, membrane resistance R_Membrane_, and cell capacitance C_cell_) are shown only for the whole-cell configuration. The intermediate cell-attached configuration (gigaseal formed but membrane patch unruptured) is shown on the bottom. Created with BioRender.

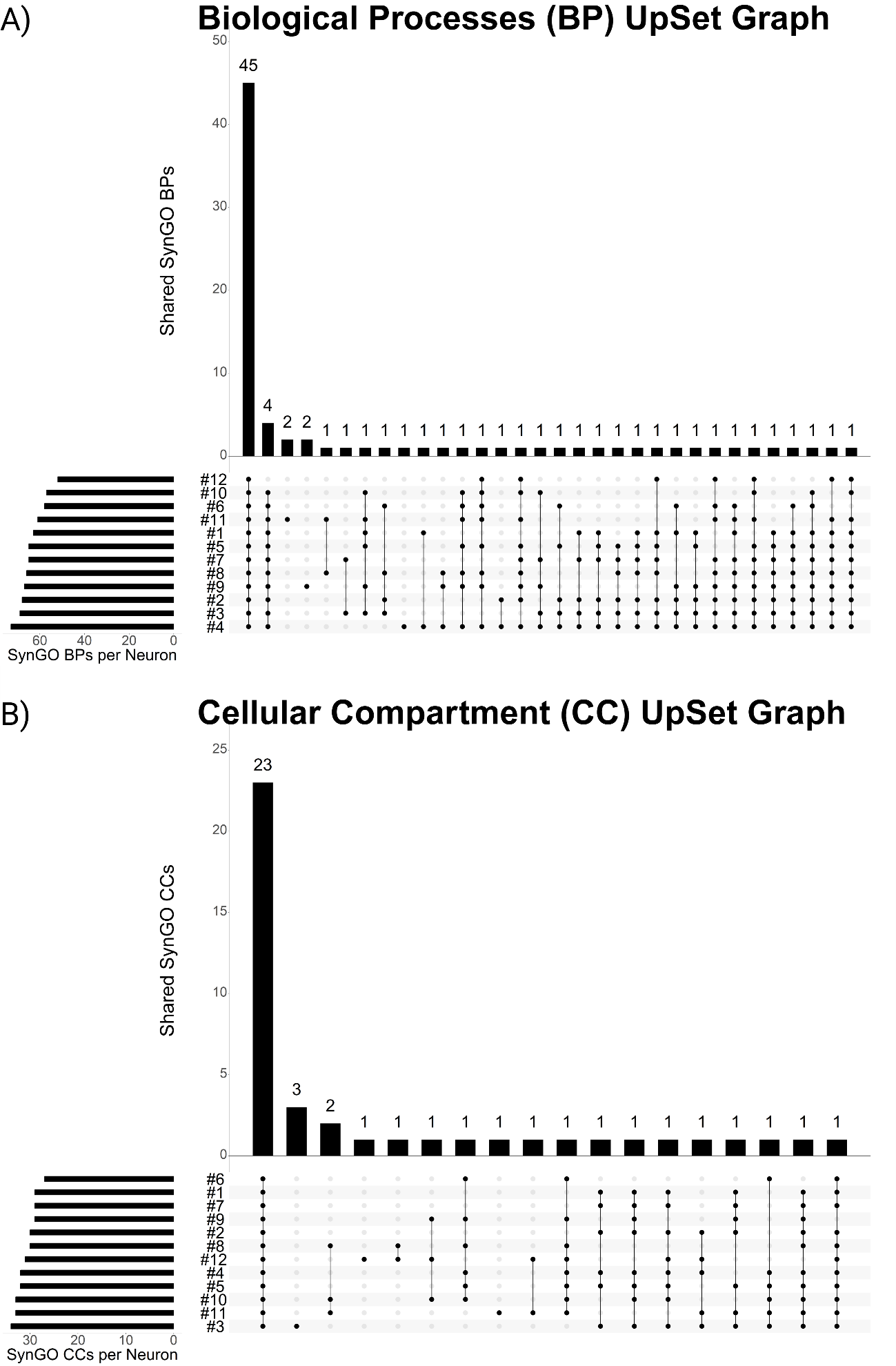

**Figure S2.** **SynGO enrichment across single neurons.** (A) biological processes (BP) and (B) cellular component (CC) gene ontology (GO) terms across single-cell samples. A core set of 45 BP and 23 CC terms significantly enriched in all 12 neurons, indicating reproducible detection of synaptic proteins. GSEA Q-value significance cutoff < 0.05

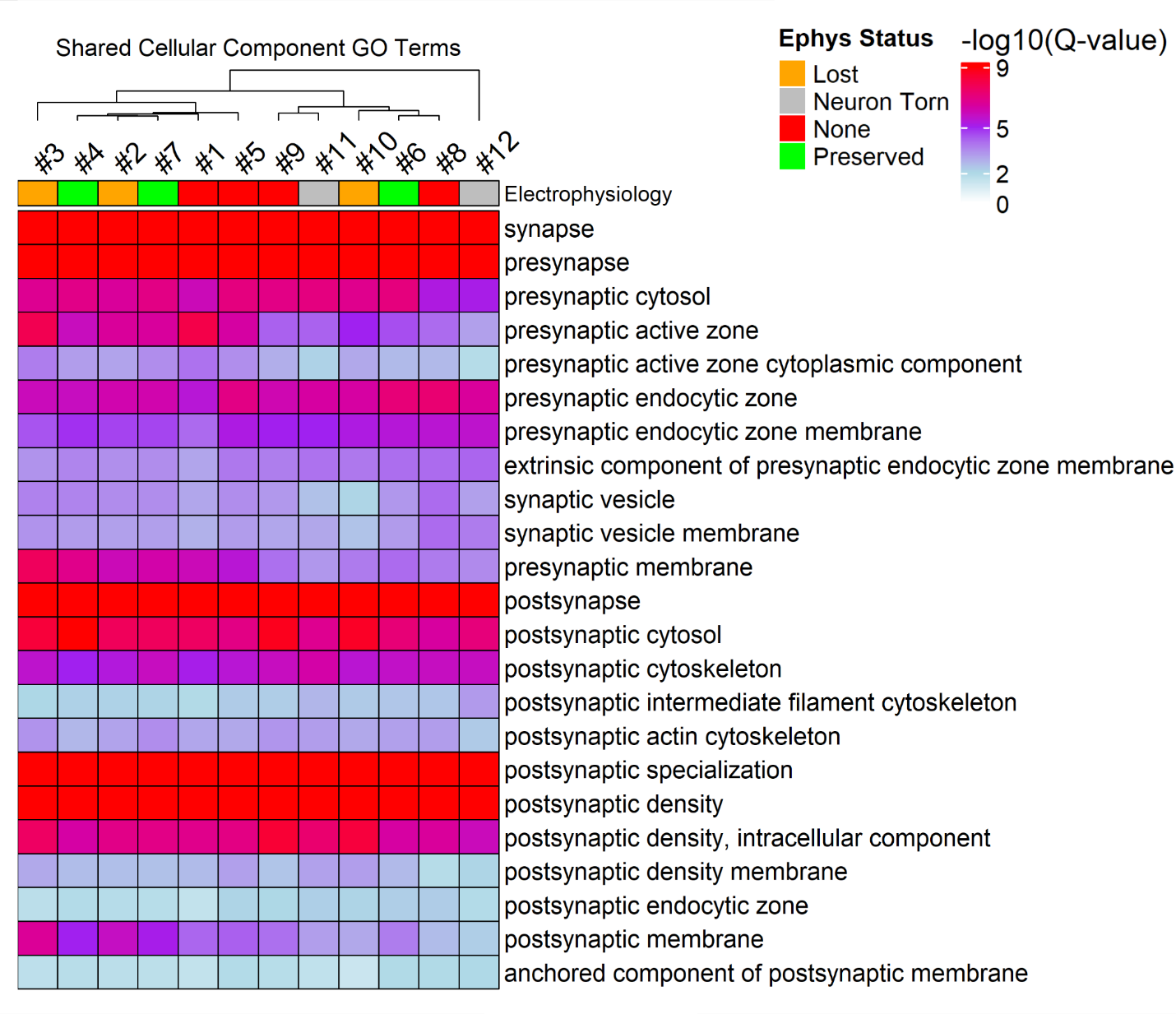

**Figure S3.** **Shared SynGO CC enrichment across single neurons.** Heatmap of 20 SynGO cellular component (CC) gene ontology (GO) terms shared across single-cell samples. Clustering was performed by samples (columns) only. GSEA Q-value significance cutoff < 0.05

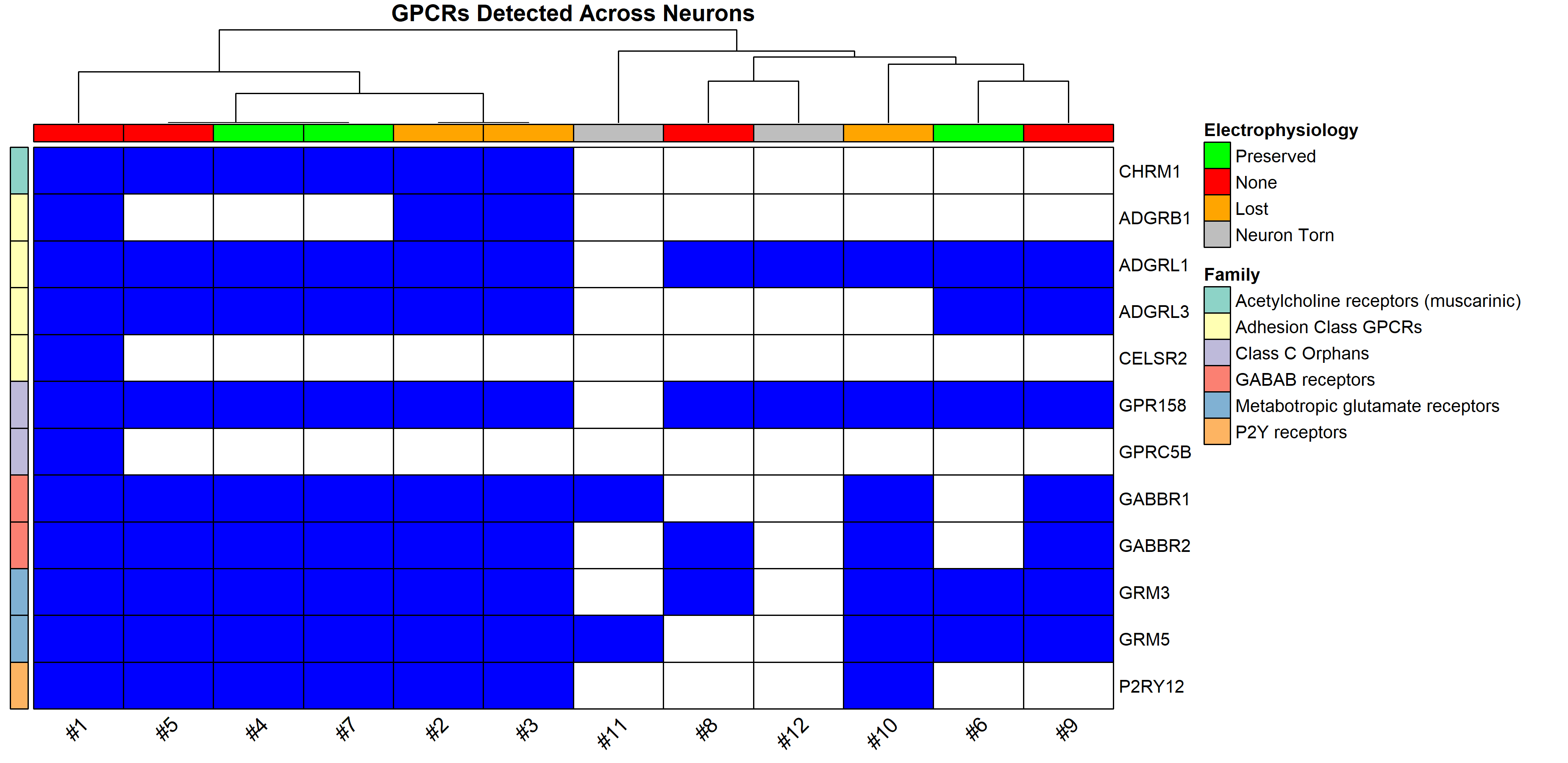

**Figure S4.** **GPCR detection across single neurons.** Binary heatmap showing presence (blue) or absence (white) of G protein-coupled receptors (GPCR) identified in individual neurons. Samples are annotated by the level of electrophysiological characterization (top bar). Green denotes whole-cell configuration was “Preserved”. Orange denotes whole-cell configuration was “Lost” during retrieval. Gray denotes whole-cell configuration was lost due to the neuron being “Torn” during retrieval. Red denotes no electrophysiological characterization due to a failure to form a gigaseal during the initial patch attempt. Clustering was performed by samples (columns) only.

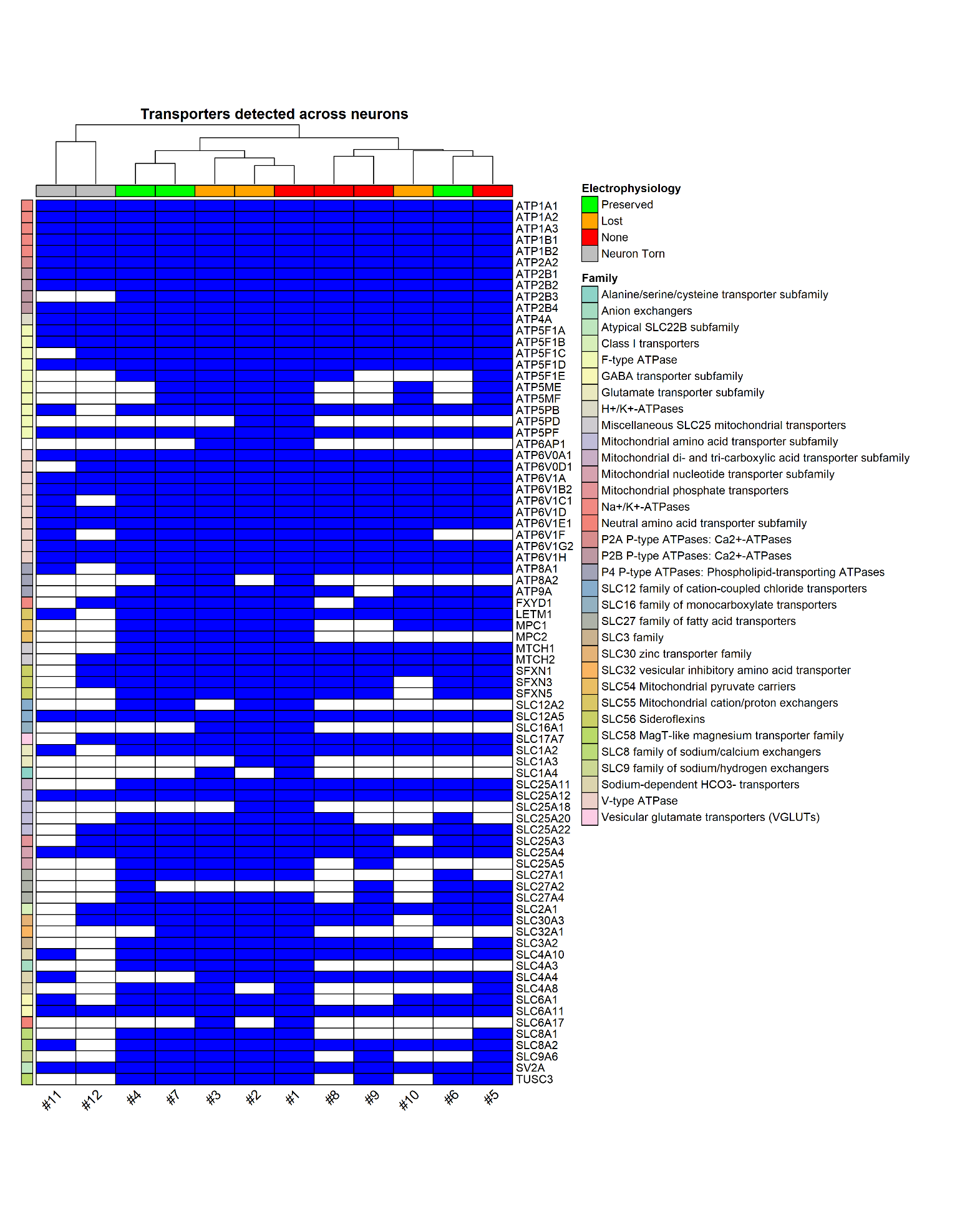

**Figure S5.** **Transporter detection across single neurons.** Binary heatmap showing presence (blue) or absence (white) of protein transporters identified in individual neurons. Samples are annotated by the level of electrophysiological characterization (top bar). Green denotes whole-cell configuration was “Preserved”. Orange denotes whole-cell configuration was “Lost” during retrieval. Gray denotes whole-cell configuration was lost due to the neuron being “Torn” during retrieval. Red denotes no electrophysiological characterization due to a failure to form a gigaseal during the initial patch attempt. Clustering was performed by samples (columns) only.
